# RASSF1A independence and early Galectin-1 upregulation in PIK3CA induced hepatocarcinogenesis: new therapeutic venues

**DOI:** 10.1101/2021.06.15.448477

**Authors:** Alexander Scheiter, Katja Evert, Lucas Reibenspies, Antonio Cigliano, Katharina Annweiler, Karolina Müller, Laura-Maria-Giovanna Pöhmerer, Timo Itzel, Silvia Materna-Reichelt, Andrea Coluccio, Kamran Honarnejad, Andreas Teufel, Christoph Brochhausen, Frank Dombrowski, Xin Chen, Matthias Evert, Diego F. Calvisi, Kirsten Utpatel

**Author notes:** **Corresponding author:** University of Regensburg, Franz-Josef-Strauß-Allee 11, 93053 Regensburg, Germany.

## Abstract

Aberrant activation of the PI3K/AKT/mTOR and Ras/Mitogen-Activated Protein Kinase pathways is a hepatocarcinogenesis hallmark. In a subset of hepatocellular carcinomas (HCC), PI3K/AKT/mTOR signaling dysregulation depends on PIK3CA mutations, while RAS/MAPK activation is partly attributed to promoter methylation of the tumor suppressor *RASSF1A*. To evaluate a possible co-carcinogenic effect of PIK3CA activation and *RASSF1A* knockout, plasmids expressing oncogenic forms of PIK3CA (E545K or H1047R mutants) were delivered to the liver of RASSF1A knockout and wildtype mice by hydrodynamic tail vein injection combined with Sleeping Beauty–mediated somatic integration. Transfection of either PIK3CA E545K or H1047R mutants sufficed to induce hepatocellular carcinomas in mice irrespective of *RASSF1A* mutational background. The related tumors displayed a lipogenic phenotype with upregulation of Fatty acid synthase and Stearoyl-CoA desaturase-1 (SCD1). Galectin-1, which was commonly upregulated in preneoplastic lesions and tumors, emerged as a regulator of SCD1. Co-inhibitory treatment with PIK3CA inhibitors and the Galectin-1 inhibitor OTX-008 resulted in synergistic cytotoxicity in human HCC cell lines, suggesting novel therapeutic venues.

**Graphical Abstract:** Hydrodynamic tail vein injection of Phosphatidylinositol-4,5-bisphosphate 3-kinase, catalytic subunit alpha (PIK3CA) mutant forms E545K and H1047R induces stepwise hepatocarcinogenesis in mice, independent of Ras association domain-containing protein 1 (RASSF1A) status. Gene expression analyses revealed an early increase in Galectin-1, which regulates the lipogenic enzyme Stearoyl-CoA desaturase-1 (SCD1). PIK3CA- and Galectin1 inhibitors act synergistically, pointing at novel therapeutic strategies.

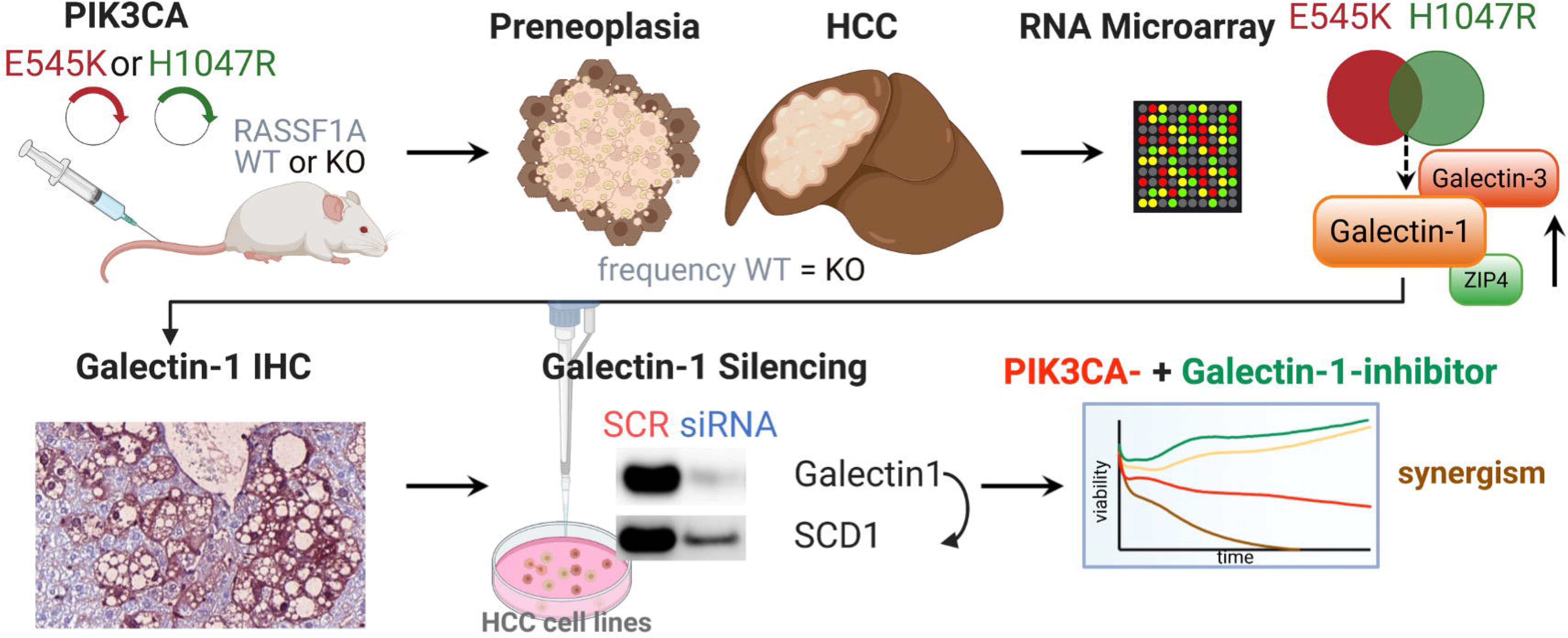

## 1. Introduction

Hepatocellular carcinoma (HCC) is the most frequent primary malignant tumor of the liver and a leading contributor to cancer mortality worldwide [1]. North America and Europe experienced a constant rising incidence of HCC over the last decades [2]. Systemic therapeutic options for advanced disease stages are scarce, albeit the combination treatment of atezolizumab and bevacizumab is one recent glimmer of hope [3]. The pathogenesis of HCC is a multistep process involving the progression from cirrhosis to low-grade dysplastic nodule, high-grade dysplastic nodule, and, ultimately, hepatocellular carcinoma [4,5]. The precise molecular mechanisms and their specific contributions to hepatocarcinogenesis remain poorly understood. Ras/MAPK signaling is among the heavily implicated pathways in HCC, despite the absence of activating *RAS* mutations [6,7]. Instead, diminished activity of Ras antagonists represents a plausible explanation for Ras/MAPK-pathway unconstrained activity [8–10]. A critical physiological counterpart of Ras is RASSF1a, which is an established tumor suppressor embedded into an intricate regulatory network [11]. RASSF1A gained importance as it is frequently epigenetically silenced by aberrant hypermethylation in diverse types of cancer [12], including HCC [6,13]. In addition to being a crucial negative-feedback downstream effector of Ras Signaling [14], RASSF1A is a regulator of microtubule stability, Rho GTPases, and the Hippo pathway [15–17]. Thereby, multiple functions are exerted, such as cell cycle arrest, inhibition of migration and metastasis through microtubular stabilization, reduction of epithelial-mesenchymal transition, and induction of apoptosis [18–20]. Concerning HCC, transfection of RASSF1A yielded growth retardation in cell lines and the respective murine xenografts [21]. Furthermore, homozygous deletion of RASSF1A elicited late liver tumor susceptibility [22] and accelerated diethylnitrosamine-induced HCC formation via autophagy promotion [23]. This leaves little doubt on the oncogenicity of RASSF1A knockout. However, the magnitude of the observed effects in the latter experimental models was somewhat limited, so that concurrent oncogenic stimuli are necessary together with RASSF1A loss to induce hepatocarcinogenesis.

Activation of the PI3K/AKT/mTOR pathway could well qualify as a promising therapeutic candidate. Indeed, the phosphoinositide 3-kinase (PI3K) cascade is one of the most deregulated pathways along tumorigenesis [14]. PI3K is a heterodimeric lipid kinase composed of a catalytic and a regulatory subunit (p110α or β or δ and p85α or β, respectively, for class Ia PI3K), which generates phosphatidylinositol (3,4,5)-trisphosphate (PIP_3_) [24]. A pivotal downstream effector is RAC-alpha serine/threonine-protein kinase (AKT), which binds to PIP3 and is subsequently activated by phosphorylation. Once induced, AKT mediates various targets and regulates numerous cellular functions, such as cell cycle progression, cell growth, survival, and metabolism [25,26]. Several genetic events triggering PI3K/AKT/mTOR activation have been reported [27]. Among them, activating mutations of the PI3K 110α subunit (PIK3CA) occupy a prominent position [28]. Important hotspot mutations include *PIK3CA* E542K, E545K, and H1047R, where the former affect the protein’s helical domain and the latter its kinase domain [29–31]. The reported PIK3CA mutational frequency in human HCCs ranges from 4 to 6 % [7,32]. Murine models have only demonstrated a hepatic co-carcinogenic effect for activated PIK3CA in conjunction with either activated yes-associated protein 1 (YAP), RasV12, or c-Met to date [33,34]. Indeed, injection of PIK3CA mutant forms alone promoted hepatic steatosis but was insufficient to induce hepatocarcinogenesis [34].

Here, we investigated a possible interplay between RASSF1A loss and the PI3K/AKT/mTOR pathway using an animal model of hydrodynamic tail vein injection with Sleeping Beauty-mediated somatic integration. E545K and H1047R PIK3CA mutant forms were injected into homozygous *RASSF1A* knockout mice. The resulting neoplastic lesions were subjected to a gene expression microarray analysis to shed light on PIK3CA driven hepatocarcinogenesis and identify putative therapeutic targets.

## 2. Materials and Methods

### 2.1. Constructs and reagents

The constructs we applied in this study comprised pCMV/sleeping beauty transposase, pT3-EF1α-PIK3CA E545K, and pT3-EF1α-PIK3CA H1047R (including human PIK3CA clones), whose generation has been described previously [33]. pT3-EF1α was used as empty vector (EV) control. Purification of plasmids was achieved using the Endotoxin-free Maxi prep kit (Sigma-Aldrich, St.Louis, MO), after which the constructs were injected into the mice.

### 2.2. Mouse breeding and genotyping

A *RASSF1A* KO founder breeding pair was kindly provided by Dr. Louise van der Weyden (Wellcome Trust Sanger Institute, Research Support Facility, Hinxton, Cambridge, CB10 1SA, UK). The genetic background of *RASSF1A* WT and KO mice was C57BL/6J x 129Sv. *RASSF1A* WT mice were obtained by crossing *RASSF1A* KO mice with C57BL/6J mice purchased from Charles River Laboratories (Sandhofer Weg, 97633 Sulzfeld, Germany). Genotyping was performed on tail biopsies after DNA extraction. Polymerase chain reactions to detect the *RASSF1A* gene were carried out according to a previously established protocol [35]. The forward primer RSF-5 (5’-CTC GCC CCT GTC AGA CCT CAA TTT CCC-3’) was applied together with the reverse primer RSF-3 (5’-CCA GGC TTC CTT CTC ACT CCT CTG CCG C3’), which yields a 400 base pair product in *RASSF1A* KO mice, where Exon 1 α has been deleted. The product length was evaluated by gel electrophoresis.

All experimental mice were kept and bred under standard conditions and stored in type III Makrolon cages with 12 h light/dark cycles. The maximum number of mice was limited to five per cage. The animals were fed autoclaved food and water ad libitum.

### 2.3. Hydrodynamic injections, mouse monitoring, and tissue sampling

Male mice of 6-8 weeks were subjected to hydrodynamic tail vein injections in any of the following groups: untreated, 1x phosphate-buffered saline (PBS), EV; pT3-EF1α-PIK3CA E545K, or pT3-EF1α-PIK3CA H1047R. Hydrodynamic tail vein injections were performed as described elsewhere [36]. Briefly, 10 µg of pT3-EF1α-PIK3CA, pT3-EF1α-PIK3CA H1047R, or pT3-EF1α were combined with pCMV/sleeping beauty transposase in a ratio of 25 to 1 in a total volume of 2 ml 0.9% sodium chloride. The solution was filtered with 0.22 µm mesh size. The total volume was injected into the lateral tail vein within 5 to 7 seconds. To attain at least 6 to 8 evaluable mice per time point and group, 10 animals per time point were injected in the experimental groups. Since control groups could be subjected to a combined analysis, only 5 animals were injected per condition and timepoint here (Table 1).

**Table 1:** Overview of experimental groups and timepoints

|  | 1 | 2 | 3 | 4 | 5 | 6 | 7 | 8 | 9 | 10 |
| --- | --- | --- | --- | --- | --- | --- | --- | --- | --- | --- |
|  | Experimental groups |  |  |  | Control groups |  |  |  |  |  |
| Strain<br>(*RASSF1A*-KO or WT) | KO | KO | WT | WT | KO | WT | KO | WT | KO | WT |
| Injected construct | H1047R | E545K | H1047R | E545K |  |  |  |  |  |  |
| 1x PBS injection |  |  |  |  | X | X |  |  |  |  |
| EV |  |  |  |  |  |  | X | X |  |  |
| untreated |  |  |  |  |  |  |  |  | X | X |
| Animals per timepoint (N) |  |  |  |  |  |  |  |  |  |  |
| 1 week | 10 | 10 | 10 | 10 | 5 | 5 | 5 | 5 | 5 | 5 |
| 1 month | 10 | 10 | 10 | 10 | 5 | 5 | 5 | 5 | 5 | 5 |
| 3 months | 10 | 10 | 10 | 10 | 5 | 5 | 5 | 5 | 5 | 5 |
| 6 months | 10 | 10 | 10 | 10 | 5 | 5 | 5 | 5 | 5 | 5 |
| 9 months | 10 | 10 | 10 | 10 | 5 | 5 | 5 | 5 | 5 | 5 |
| 12 months | 10 | 10 | 10 | 10 | 5 | 5 | 5 | 5 | 5 | 5 |
| Sum of animals per experimental group | 60 | 60 | 60 | 60 | 30 | 30 | 30 | 30 | 30 | 30 |
| Sum of all animals | 420 |  |  |  |  |  |  |  |  |  |

Animals were excluded from the evaluation if the injected total volume was below 2 ml. Another criterium to assess the quality of injection was the mouse behavior following the injection. Successfully injected mice displayed a decreased activity for at least 60 minutes due to the systemic volume challenge. Mice that did not show a similar behavior were excluded from the experiment. Mice were monitored daily and kept until the prespecified experimental time point. Respiratory distress, lethargy, and palpable liver masses equivalent to a size of ∼3.5 to 4 cm were defined as termination criteria. The cervical dislocation was applied. A photo-documentation of the livers was performed, after which half of the livers was frozen in liquid nitrogen, and the other half was formalin-fixed and paraffin-embedded. 1-2 mm^3^ of liver tissue were fixed in glutaraldehyde. Animal breeding and all conducted animal experiments were in accordance with protocols by the Mecklenburg-Western Pomeranian federal institution “Landesamt für Landwirtschaft, Lebensmittelsicherheit und Fischerei (LALLF) Mecklenburg-Vorpommern” (protocol number/Aktenzeichen: 7221.3-1.1-052/12).

### 2.4. Histology, immunohistochemistry, image acquisition, and proliferation index

At least two board-certified pathologists and liver experts conducted the histopathological assessment of the liver lesions (K.U., K.E.). The liver tissue was processed following the standard diagnostic institutional guidelines. Two μm-thin histological sections were cut from formalin-fixed, paraffin-embedded tissue samples. The sections were deparaffinized through a series of xylene and gradient alcohols to water. For antigen retrieval, the slides were heated in a microwave oven for 10 minutes, while 10 mM sodium citrate buffer with a pH of 6.0 was applied. Afterward, the samples were cooled down to room temperature. The slides were treated with 1× Dako Peroxidase-Blocking Solution® (cat S2023; Agilent Technologies Inc., Santa Clara, CA) for 10 min. The primary antibody was diluted and applied in Dako Antibody Diluent® (cat. S2022, Agilent Technologies Inc.); the slides were placed in a humidity chamber and incubated at room temperature overnight (see Table 2 for a list of all primary antibodies). Following a subsequent washing step with Dako washing solution® (cat. S3006, Agilent Technologies Inc.), the secondary antibody Histofine Simple Stain MAX PO® anti-goat or anti-rabbit (Nacalai USA, Inc., San Diego, CA) was administered for 60 min at room temperature. Two more washes in Dako washing solution® followed. For chromogenic reactions, we employed Dako Liquid DAB + Substrate Chromogen System® (cat. K346811-2, Agilent Technologies Inc.), according to the manufacturer’s instructions. Counterstaining was performed with Mayer’s hemalum for 10 s. The application of coverslips was made automatically using Ventana BenchMark Ultra® (Roche, Penzberg, Germany).

**Table 2:**
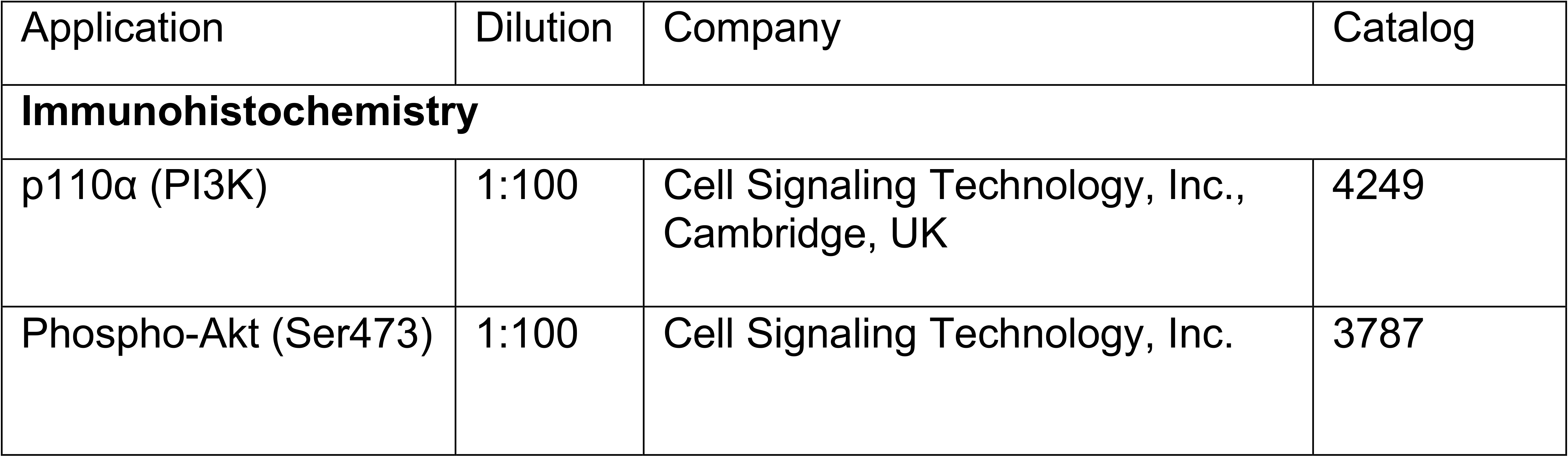
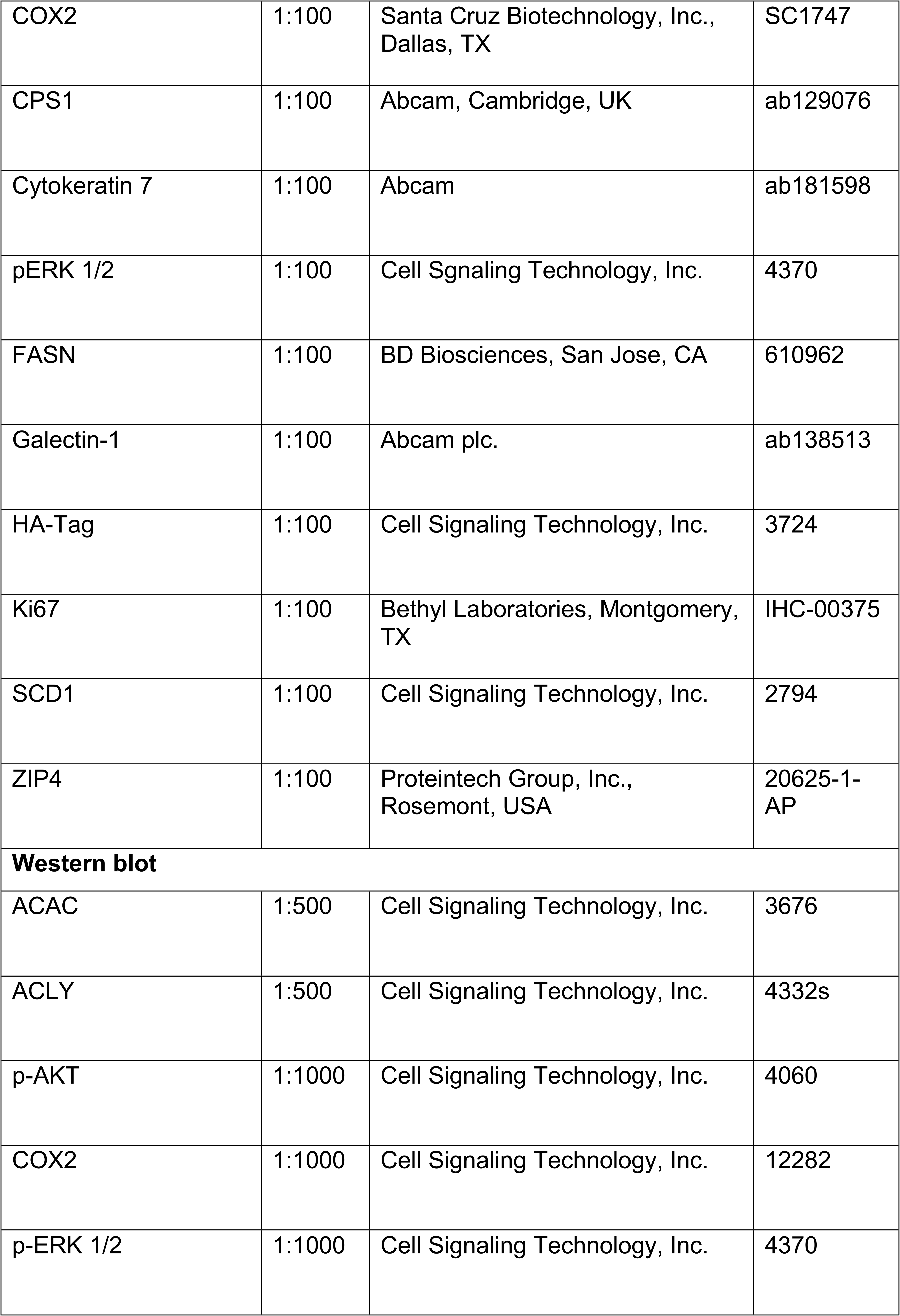
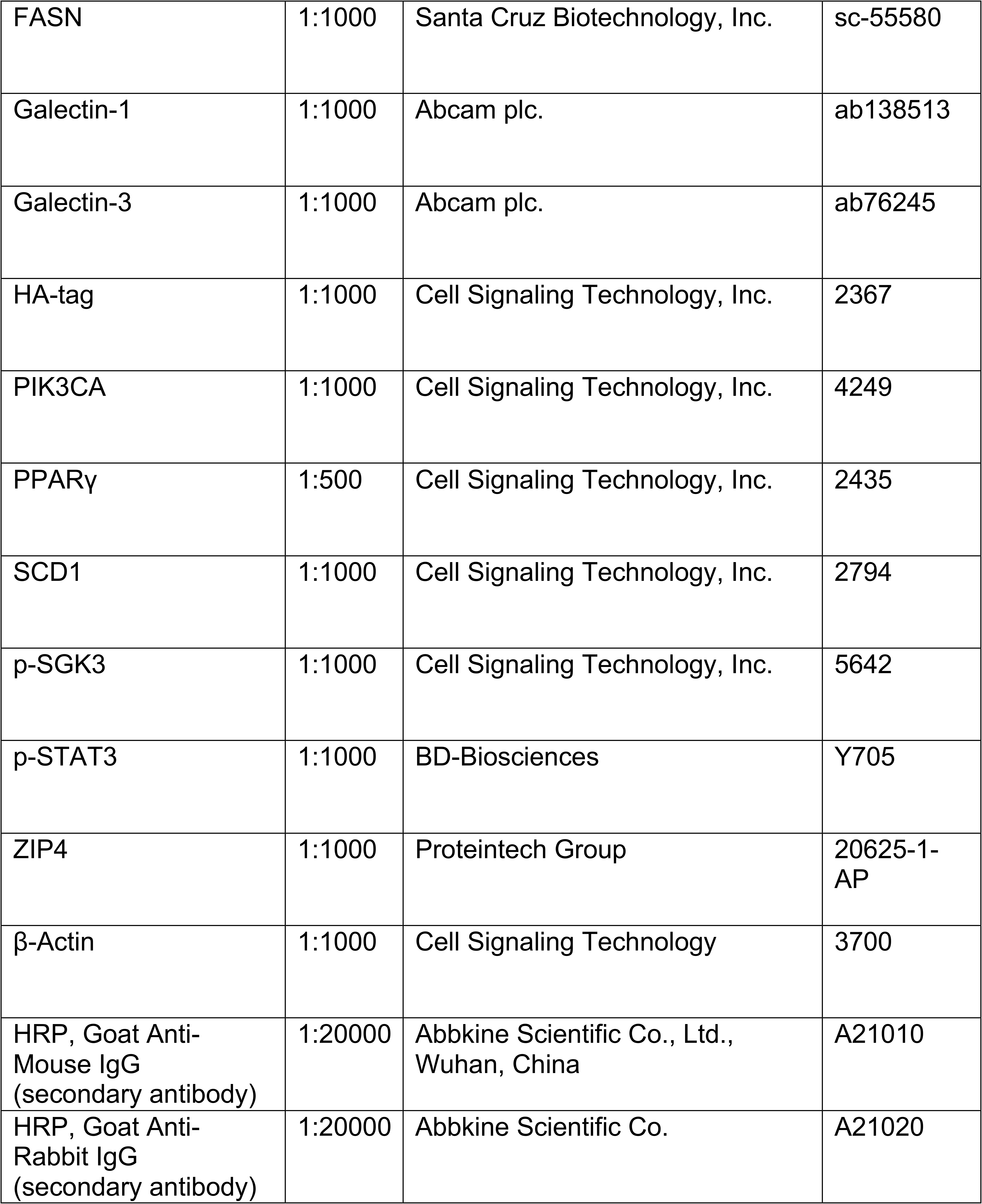
Antibodies used for immunohistochemistry and Western blots

| Application | Dilution | Company | Catalog |
| --- | --- | --- | --- |
| **Immunohistochemistry** |  |  |  |
| p110α (PI3K) | 1:100 | Cell Signaling Technology, Inc., Cambridge, UK | 4249 |
| Phospho-Akt (Ser473) | 1:100 | Cell Signaling Technology, Inc. | 3787 |
| COX2 | 1:100 | Santa Cruz Biotechnology, Inc.,<br>Dallas, TX | SC1747 |
| CPS1 | 1:100 | Abcam, Cambridge, UK | ab129076 |
| Cytokeratin 7 | 1:100 | Abcam | ab181598 |
| pERK 1/2 | 1:100 | Cell Signaling Technology, Inc. | 4370 |
| FASN | 1:100 | BD Biosciences, San Jose, CA | 610962 |
| Galectin-1 | 1:100 | Abcam plc. | ab138513 |
| HA-Tag | 1:100 | Cell Signaling Technology, Inc. | 3724 |
| Ki67 | 1:100 | Bethyl Laboratories, Montgomery,<br>TX | IHC-00375 |
| SCD1 | 1:100 | Cell Signaling Technology, Inc. | 2794 |
| ZIP4 | 1:100 | Proteintech Group, Inc.,<br>Rosemont, USA | 20625-1-AP |
| **Western blot** |  |  |  |
| ACAC | 1:500 | Cell Signaling Technology, Inc. | 3676 |
| ACLY | 1:500 | Cell Signaling Technology, Inc. | 4332s |
| p-AKT | 1:1000 | Cell Signaling Technology, Inc. | 4060 |
| COX2 | 1:1000 | Cell Signaling Technology, Inc. | 12282 |
| p-ERK 1/2 | 1:1000 | Cell Signaling Technology, Inc. | 4370 |
| FASN | 1:1000 | Santa Cruz Biotechnology, Inc. | sc-55580 |
| Galectin-1 | 1:1000 | Abcam plc. | ab138513 |
| Galectin-3 | 1:1000 | Abcam plc. | ab76245 |
| HA-tag | 1:1000 | Cell Signaling Technology, Inc. | 2367 |
| PIK3CA | 1:1000 | Cell Signaling Technology, Inc. | 4249 |
| PPARγ | 1:500 | Cell Signaling Technology, Inc. | 2435 |
| SCD1 | 1:1000 | Cell Signaling Technology, Inc. | 2794 |
| p-SGK3 | 1:1000 | Cell Signaling Technology, Inc. | 5642 |
| p-STAT3 | 1:1000 | BD-Biosciences | Y705 |
| ZIP4 | 1:1000 | Proteintech Group | 20625-1-AP |
| β-Actin | 1:1000 | Cell Signaling Technology | 3700 |
| HRP, Goat Anti-Mouse IgG (secondary antibody) | 1:20000 | Abbkine Scientific Co., Ltd., Wuhan, China | A21010 |
| HRP, Goat Anti-Rabbit IgG (secondary antibody) | 1:20000 | Abbkine Scientific Co. | A21020 |

Images were acquired with the slide scanner Pannoramic 250 Flash III® (Sysmex, Kobe, Japan) with a 20x objective. Subsequently, images were displayed after stitching using the software CaseViewer® (Sysmex). With 300 pixels per inch, resolution screenshots were taken.

The stitched images of Ki67 immunohistochemistry were imported to the software DeePathology^TM^ STUDIO (DeePathology.ai; Raanana, Israel). A board-certified pathologist (K.U.) trained the algorithm to discern hepatocyte nuclei from nuclei of other cell types. These were determined as background cells (such as lymphocytes and Kupffer cells) and excluded from the analysis. Regions of interest were manually selected. A graphical representation of the calculation results was visually assessed for further validation.

### 2.5. Western blot analysis

For protein extraction, lysis of cells and mouse liver tissues was achieved using the Mammalian Protein Extraction Reagent (cat 78501; Thermo Fisher Scientific, Waltham, MA) in association with the Halt Protease Inhibitor Cocktail (cat 78429; Thermo Fisher Scientific). An additional mechanical force was applied for tissue homogenization with the Next Advance Bullet Blender® Storm 24 (Next Advance, Inc.; Troy, NY). The lysates were incubated for 30 minutes at 4°C while vortexing every 5-10 minutes. A centrifugation step for 30 minutes (>20,000 g, 4°C) ensued. Concentrations of total protein were assessed using the Bradford Protein Assay [37]. To attain a standard curve for linear regression, serial dilutions of BSA were employed.

For Western blot analysis, protein lysates were separated by sodium dodecyl sulfate-polyacrylamide gel electrophoresis. 2.5 μg of total proteins were loaded onto BoltTM 4-12% Bis-Tris Plus gels (Thermo Fisher Inc.) at 150 V for 30-60 min. Next, gels were incubated in 10% ethanol for dehydration before blotting the proteins to a membrane using the BlotTM 2 Gel Transfer Device (Thermo Fisher Inc.). After staining the membranes in Ponceau solution, they were placed in EveryBlot Blocking Buffer (Bio-Rad Laboratories, Inc.; Hercules, CA) for 15-30 minutes at room temperature. Primary antibodies (Table 2) were diluted in the blocking buffer and incubated overnight at 4°C. The next day, membranes were washed in Tris Buffered Saline with Tween® (Cell Signaling Technology, Inc.) for 5 minutes at room temperature. Then, the secondary antibody was applied at room temperature for 1 hour. The membranes were washed with Tris Buffered Saline with Tween®. The chemifluorescent signal was visualized by Clarity Max^TM^ Western ECL Substrate (Bio-Rad Laboratories) on a ChemiDoc^TM^ MP Imaging System (Bio-Rad Laboratories). For quantitative analysis of band intensities, version 6.1 of the software ImageLab (Bio-Rad Laboratories) was employed. Values were normalized to the corresponding β-actin bands.

### 2.6. Transmission electron microscopy

Liver tissue samples were fixed in 0.1 M cacodylate-buffered Karnovsky fixative containing 2.5% glutaraldehyde and 2% paraformaldehyde (Electron Microscopy Sciences, Hatfield, PA) overnight at room temperature with a subsequent post-fixation in 1% osmium tetroxide (Electron Microscopy Sciences), which was applied for 2 h. Next, the samples were dehydrated in graded ethanols (Sigma Aldrich). Afterward, they were embedded in an EMbed-812 epoxy resin (Electron Microscopy Sciences). Following two days of heat polymerization at a temperature of 60°C, 0.8 µm thin sections were prepared. These were stained with toluidine blue (AgarScientific; Essex, United Kingdom) and basic fuchsine solution (Polysciences Inc.; Warrington, PA). Subsequently, the epon block was adjusted to allow ultrathin sectioning. Eighty nm sections were cut with a diamond knife on a Reichert Ultracut-S ultramicrotome (Leica, Wetzlar, Germany). These were double contrasted using aqueous 2% uranyl acetate (Honeywell International Inc.; Morristown) and lead citrate solutions (Leica) for 10 min each. A LEO912AB transmission electron microscope (Zeiss, Oberkochen, Germany) operated at 100 kV was used for imaging the ultrathin sections.

### 2.7. Gene expression microarray hybridization and analysis

RNA was isolated from liver tissue using the NucleoSpin® RNA Plus Kit (Macherey-Nagel GmbH & Co. KG, Düren, Germany) and following the manufacturer’s instructions. Gene expression microarray analysis was performed with mouse Sentrix® BeadChips (Illumina; San Diego, CA). We followed the workflow established by the manufacturer. Briefly, cDNA synthesis, in vitro transcription, and cleanup were conducted using the Ambion® RNA Amplification Kit (Illumina). The samples were quantified after the addition of Molecular Probes Ribo Green® (Thermo Fisher Scientific) by fluorometric measurement. Afterward, the samples were hybridized, and an image was extracted with the BeadArray Reader (Illumina). The data were analyzed with the BeadStudio® application (Illumina).

Raw data were background corrected, quantile-normalized, and log2-transformed by using the limma package [38] provided by R/Bioconductor (https://www.bioconductor.org). Subsequently, differential expressed genes where calculated, again using the limma package. Concerning multiple testing, Benjamini and Hochberg’s method was applied to adjust the p-values. Heatmaps were generated with the pheatmap package provided by R/Bioconductor (https://www.bioconductor.org). Furthermore, pre-ranked Gene Set Enrichment Analysis (GSEA) (Hallmark gene sets) were performed using the official software tools from the Broad Institute (Boston, MA, USA, https://www.gsea-msigdb.org) [39]. For this purpose, all probes representing the same gene symbol were averaged.

### 2.8. Cell culture and *in vitro* studies

The human HCC cell lines PLC/PRF/5, HLE, HLF, and Snu182 were cultured in 5% CO_2_ at 37°C in a humidified incubator. Cells were grown in Dulbecco’s modified Eagle medium (Gibco, Grand Island, NY) or RPMI 1640 Medium (Gibco) supplemented with 5% fetal bovine serum (Gibco), 100 mg/mL streptomycin, and 100 U/mL penicillin.

We performed cell viability assays using the xCELLigence® real-time cell analysis dual plate (RTCA DP) device (OLS OMNI Life Science GmbH & Co KG; Bremen, Germany). For impedance-based real-time cell index measurement, cells were grown on E-Plate 16 PET (Agilent Technologies, Inc.). Measurement sweeps were acquired every 15 minutes. Six thousand two hundred fifty cells suspended in a total volume of 150 µl of growth medium were seeded in each well. After 24 hours, varying concentrations of Dimethyl sulfoxide (Sigma-Aldrich), the inhibitory compounds Alpelisib (MedChemExpress; LLC., Monmouth, NJ), and/or OTX008 (MedChemExpress) were added. Afterward, the measurement was acquired for a total of 72 hours. Raw data were analyzed with the RTCA software (OLS OMNI Life Science GmbH & Co KG). The data were normalized to the timepoint of inhibitor addition. Synergism was evaluated 24 hours after adding the inhibitory compounds using the software CompuSyn (ComboSyn, Inc., Paramus, NJ), which generates dose-effect curves and isobolograms, and determines the combination index.

For Alpelisib single treatment, cells were seeded in 6 well plates at a density of 3-5 x 10^5^ cells in 2 ml medium per well. The next day, either Alpelisib in varying concentrations or matched DMSO was added. After an incubation of 48 hours, cells were harvested, centrifuged and the cell pellet was used for further processing.

For Galectin-1 silencing, cells were seeded at a density of 3×10^5^ cells in 2 ml of medium per well in 6 well plates. Cells were transfected with Silencer® Select Negative Control #1 siRNA (Thermo Fisher Inc) or LGALS1 siRNA (Eurofins Genomics; Ebersberg, Germany) with the sense sequence 5’-[UUGCUGUUGCACACGAUG-GUGUUGG]-3’ the following day using Lipofectamine^®^ RNAiMAX (Thermo Fisher Inc) according to the manufacturer’s instructions. Lipofectamine and siRNA were diluted in OptiMEM^®^ Reduced Serum Medium (Thermo Fisher Inc.) and combined. Medium in the wells was discarded, and cells were washed with 1x PBS before adding the transfection solution. After an incubation period of 48 h, the transfection was repeated once more. Cells were harvested after an additional 24 h of incubation using cell scrapers. Harvested cell suspensions were centrifuged (300g, 5 min). The pelleted cells were used for further analyses.

### 2.10. ATP detection assay and drug screening

PLC/PRF/5 cells were seeded in 384 well plates (µClear #781091, Greiner Bio-One GmbH, Frickenhausen, Germany) at a density of 2500 cells per well. Cells were treated at the time of seeding with the set concentration of OTX008 20 µM (#35318, MedChemExpress), Alpelisib (S2814, Selleck Chemicals Llc, Houston, TX) 1 µM, Buparlisib 1 µM (S2247, Selleck Chemicals Llc) or Taselisib 1 µM (S7103, Selleck Chemicals Llc) either alone or in combination using the D300e digital dispenser (Tecan Group, Männedorf, Switzerland). Cell viability was measured 72 hours after treatment using an ATPlite 1step detection assay (#6016739, PerkinElmer, Inc., Waltham, MA). To assess synergy between OTX008 and the PI3K inhibitors, we used the Bliss model of independence [40] to calculate the expected effect of the combination treatment, assuming the compounds act independently.

#### Expected_AB_=E_A_+E_B_-EA_EB_

where E_A_ and E_B_ are the effects, measured as the reduction in viability, of the two compounds in monotherapy. The observed effect of the combination therapy is directly compared to the expected effect according to the Bliss model and calculated as the excess over the Bliss score (EOB) [41].

#### EOB=Observed_AB_-Expected_AB_

If EOB is >0, the combined compounds have an effect more potent than if they acted independently.

A compound library including 315 approved anti-cancer drugs was purchased from TargetMol (L2110, Target Molecule Corp., Boston, MA), diluted to a concentration of 1 mM and 40 nL were dispensed in 384 well plates (Greiner µClear #781091, Greiner Bio-One GmbH) using an acoustic liquid handler (Echo® 550, Beckman Coulter Life Sciences, Brea, CA). PLC/PRF/5 cells were detached and separated in two tubes containing either OTX008 at a final concentration of 20 µM or DMSO vehicle. The cells were then seeded using a liquid dispenser (Multidrop™ Combi, Thermo Fisher Scientific) at 2500 cells per well in pre-spotted assay plates at a final volume of 40 µL per well so that the compounds are diluted to a final concentration of 1 µM. Cells were incubated at 37°C, 5% CO_2_ for 72 hours. At the end of the incubation time, 10 µL of a solution of PBS and Hoechst 33342 (2 drops/mL, NucBlue™ Live ReadyProbes™ Reagent, Thermo Fisher Scientific) was added to each well using a liquid dispenser (Multidrop™ Combi, Thermo Fisher Scientific), incubated for 1 hour and imaged for brightfield and Hoechst staining using the Operetta CLS™ High-Content Analysis System (PerkinElmer, Inc.). Stained nuclei were counted using the Harmony® software (PerkinElmer, Inc.), and the number of nuclei was used to measure cell viability and proliferation. For each plate, the proteasome inhibitor Carfilzomib was used as a positive control and DMSO as a negative control to calculate the Z’ value as a plate QC criteria: all plates had a Z’ > 0.7, above the widely accepted threshold of 0.5. Data from two independent experiments were averaged (correlation coefficient between replicates 0.96). The number of nuclei in each well was normalized to DMSO-treated controls, and normalized viability was used to calculate the excess over the Bliss score (EOB) as described above. Compounds were defined as a hit if the observed normalized viability upon treatment in combination with OTX008 was below 0.6 and the Z score of EOB was greater than 1.5. For each compound, information regarding the mode of action and the targeted genes was obtained from the drug repurposing hub database [42].

### 2.10. Statistical analyses

Tumor occurrence frequency was compared by descriptive statistics. Comparisons between two groups were conducted with non-parametric Mann-Whitney-U tests due to small sample sizes (quantification of lipogenic enzymes in Galectin-1 silencing experiments). For multiple comparisons, non-parametric data were compared using the Kruskal-Wallis test without adjustment (liver/weight over body weight comparison; quantification of Western blots for Galectin-1 in PIK3CA E545K liver lysates; quantification of Western Blots in Alpelisib treatment experiments).

Kaplan-Meier curves and Log-rank test were used to compare survival between *RASSF1A* WT and KO mice with PIK3CA E545K injection.

A mixed linear model (maximum likelihood estimation, unstructured repeated covariance type) was conducted to evaluate repeated measures of proliferation between *RASSF1A* KO and WT mice and assess the proliferation within normal tissue, preneoplastic lesions, and tumors. Moreover, corresponding injection groups were compared against each other separately in *RASSF1A* WT versus KO mice.

GraphPad Prism version 9, GraphPad Prism R version 4.0.3 (GraphPad Software; San Diego, CA), and SPSS version 26 (IBM; Armonk, NY) were employed. All *p* values were obtained in two-tailed tests, and *p* ≤ 0.05 was considered statistically significant. Microarray data were analyzed using the R package Limma. Moderated contrast t-tests were computed, and Benjamini and Hochberg’s method to adjust for multiple testing was applied to microarray expression data [43,44].

## 3. Results

### 3.1. PIK3CA mutant forms E545K and H1047R cause hepatocarcinogenesis irrespective of *RASSF1A* mutational background

A mouse model of hydrodynamic tail vein injection with Sleeping Beauty-mediated somatic integration [45–48] was chosen to evaluate RASSF1A and PIK3CA mutant forms carcinogenic cooperativity. This procedure favors the transfection of pericentral hepatocytes (acinus zone 3) and yields transfection efficiencies in the range of 5 to 10 % of hepatocytes (Figure 1*A*) [49].

**Figure 1.**
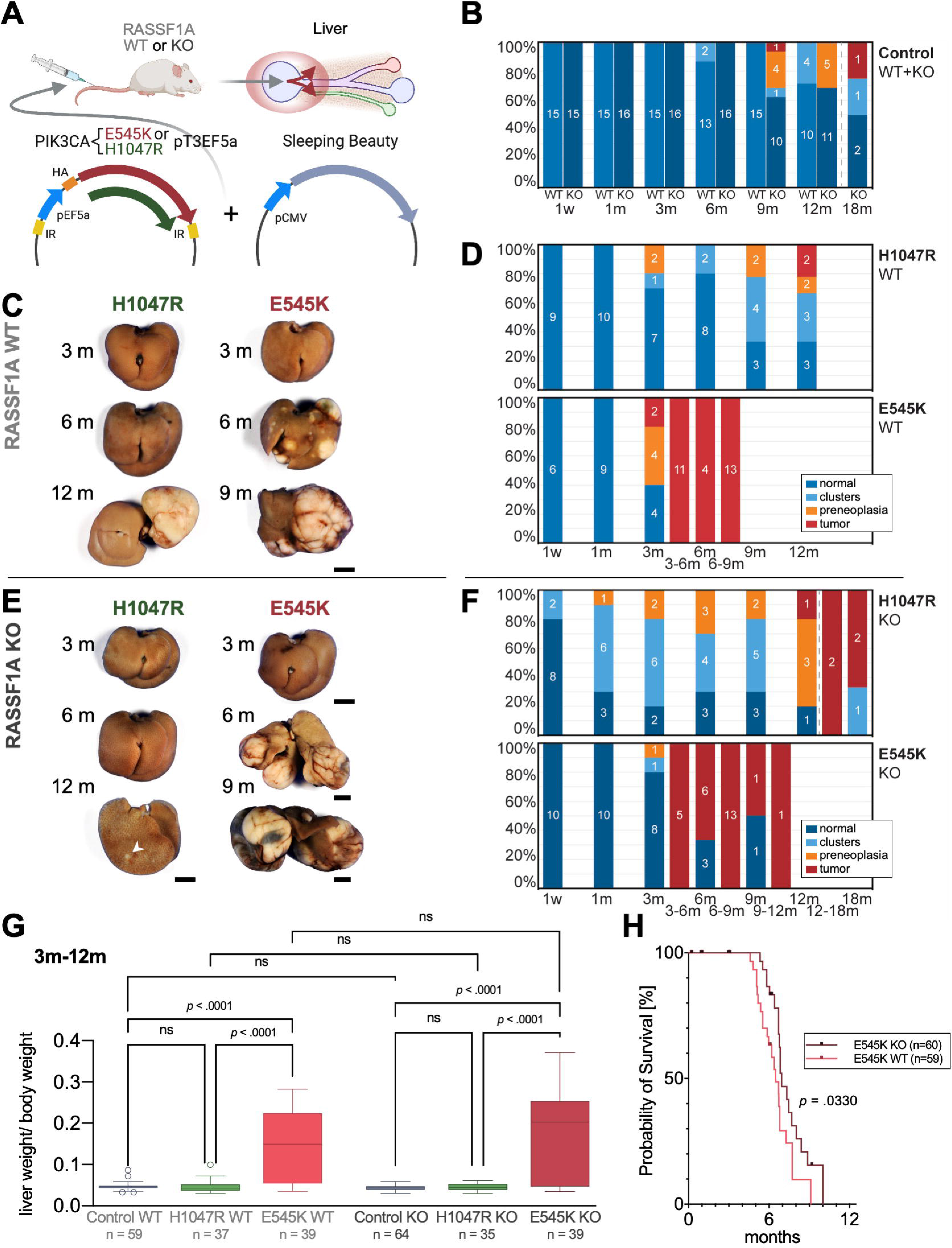
Hydrodynamic injection of PIK3CA mutated forms in the mouse liver drives hepatocarcinogenesis irrespective of RASSF1A background. *(A)* Scheme of hydrodynamic tail vein injections of PIK3CA mutant forms E545K or H1047R in conjunction with a plasmid encoding Sleeping Beauty transposase into either RASSF1As wildtype (WT) or knockout (KO) mice leading to preferential transfection of pericentral hepatocytes as indicated by the red circle and arrows in the liver scheme. *(B)* Stacked bar charts showing the occurrence frequency of clear-cell clusters, preneoplastic lesions, and tumors as well as normal-appearing liver tissue (specified in the legend) at noted time points based on histological examination in RASSF1A WT versus KO mice in combined injection control groups (untreated, PBS-injected, and empty-vector injected mice). Central digits specify the number of observed animals in the respective classification subgroups. The dashed line is indicative of an extended observation period for 4 RASSF1A KO mice. *(C)* Representative gross images of livers of RASSF1A WT mice injected with PIK3CA mutant forms H1047R and E545K at noted time points. *Scale bar:* 0.5 cm. *(D)* Stacked bar charts visualizing the frequency of occurrence of neoplastic lesions (specified in the legend) in PIK3CA H1047R and E545K injected RASSF1A WT mice. *(E)* Representative gross images of livers of RASSF1A KO mice injected with PIK3CA mutant forms H1047R and E545K at noted time points. The arrowhead points at a small tumor. *Scale bars:* 0.5 cm. *(F)* Stacked bar charts visualizing the frequency of occurrence of neoplastic lesions (specified in the legend) in PIK3CA H1047R and E545K injected RASSF1A KO mice at noted time points. *(G)* Diagram comparing the ratio of liver weight and body weight as tumor burden surrogate for combined timepoints ranging from 3 months to 12 months in the respective PIK3CA injection groups sorted by RASSF1A background. Tukey method box plots are displayed. A Kruskal-Wallis test was calculated. *(H)* Kaplan-Meier survival curves (euthanasia based on termination criteria) of RASSF1A WT and KO mice from PIK3CA E545K injection group. Log-rank test showed a minor significant longer survival time in KO than WT mice (p = .0330). Mice number in each arm is labeled in the figure legend.

Contrary to our expectations from previous injections of mutant PIK3CA containing plasmids [33,34], mice in all PIK3CA injection groups developed liver tumors within the prespecified observation period of 12 months. This unexpected finding presumably depends on the mixed background used for the experiments (C57BL/6J x 129Sv), which differs from the previously employed FVB/N inbred mouse strain [34].

Based on combined gross and histological examination, the observed lesions were stratified into the following categories: preneoplastic lipid-rich clusters, preneoplasias (criteria of expansive growth with initial compression of surrounding tissue and estimated cell content >100), and tumors (irregular borders, presence of necrosis, expansive growth with evident compression or diffuse infiltration of surrounding tissue, macroscopic correlation, cytologic signs of malignancy). Tumors in PIK3CA E545K injected mice were already detectable after 3 months, instead of a 12 months latency of tumorigenesis in the PIK3CA H1047R injection groups. H1047R injections yielded discernible tumors in 3 of 112 mice within the defined observation time of 12 months. In contrast, E545K injections resulted in numerous tumors (tumors in 56 of 113 mice), which frequently necessitated a premature termination. Serving as solid evidence against cooperativity, neither clusters, preneoplastic lesions, nor tumors displayed a marked difference in the occurrence frequency when comparing PIK3CA-mutant forms injected in *RASSF1A* wildtype (WT) and RASSF1A knockout (KO) mice. A combined control group (∼5 mice each without transfection, transfection of 1x phosphate-buffered saline, and transfection of the empty vector) did not demonstrate spontaneous tumorigenesis in *RASSF1A* WT mice. However, the *RASSF1A* knockout mice control group developed a total of two tumors at the experimental time points of 9 and 18 months (extended period of observation; Figure 1*B-F*).

To assess differences in tumor burden between experimental groups, the liver weight/body weight ratio for the combined experimental timepoints ranging from 3 months to 12 months were compared using the Kruskal-Wallis test (Figure 1*G*). The post-hoc test showed that PIK3CA E545K possesses an increased oncogenic potency compared to H1047R and control in *RASSF1A* WT and KO mice (*p values* < 0.0001). PIK3CA H1047R injected mice were neither significantly different from control in *RASSF1A* WT nor KO mice (*p values* > 0.05). Moreover, the lack of significant differences between *RASSF1A* KO and WT mice substantiated that *RASSF1A* loss does not increase tumor burden (*p values* > 0.05).

Next, we conducted a survival analysis in mice with tumor-related deaths occurring only in PIK3CA E545K subgroups. It should be noted that animals were deliberately euthanized based on the assessment of the following termination criteria: respiratory distress, lethargy, and palpable liver masses equivalent to a size of ∼3.5 to 4 cm. Premature termination took place at the formerly denoted timepoints 3-6 months, 6-9 months, and 9-12 months. A minor, longer survival of PIK3CA E545K injected mice was detected in *RASSF1A* KO mice (median survival= 6.9 months) compared to *RASSF1A* WT mice (median survival = 6.5 months, *p* = .0330) (Figure 1*H*).

Altogether, these data indicate that *RASSF1A* loss does not increase the susceptibility to PIK3CA mutant forms-mediated hepatocarcinogenesis and even shows a tendency to improve survival in the PIK3CA E545K group. Of note, the only two tumors observed in the control group both emerged in *RASSF1A* knockout mice. Even if these tumors were indeed *RASSF1A* related, a very long latency was required for tumorigenicity.

Next, we searched for potential histological differences between *RASSF1A* WT and KO mice. Histological analysis revealed that proliferative clusters, pericentrally located preneoplastic lesions, and tumors were characterized by a lipid-rich phenotype and possibly shared the common ancestor of disseminated lipid-rich cells, which can be observed in the liver acinus zone 3, corresponding to singular transfected cells. PIK3CA H1047R injected mice preferentially developed discontinuous but expansive preneoplastic lesions in the sense of confluent clusters, while PIK3CA E545K injected mice developed focal, coherent, and more rounded preneoplastic lesions. Tumors were predominantly well-differentiated and displayed a relatively low nuclear-cytoplasmic ratio and a low degree of nuclear pleomorphism while maintaining their lipid-rich phenotype. Visual examination of Ki67 revealed an increase of proliferation in tumors of E545K injected mice, which was not readily apparent for the observed lipid-rich clusters in the H1047R injection group. There was no histologically evident difference in the morphology of preneoplastic lesions and tumors between *RASSF1A* WT and KO mice (Figure 2*A* and *B*).

**Figure 2.**
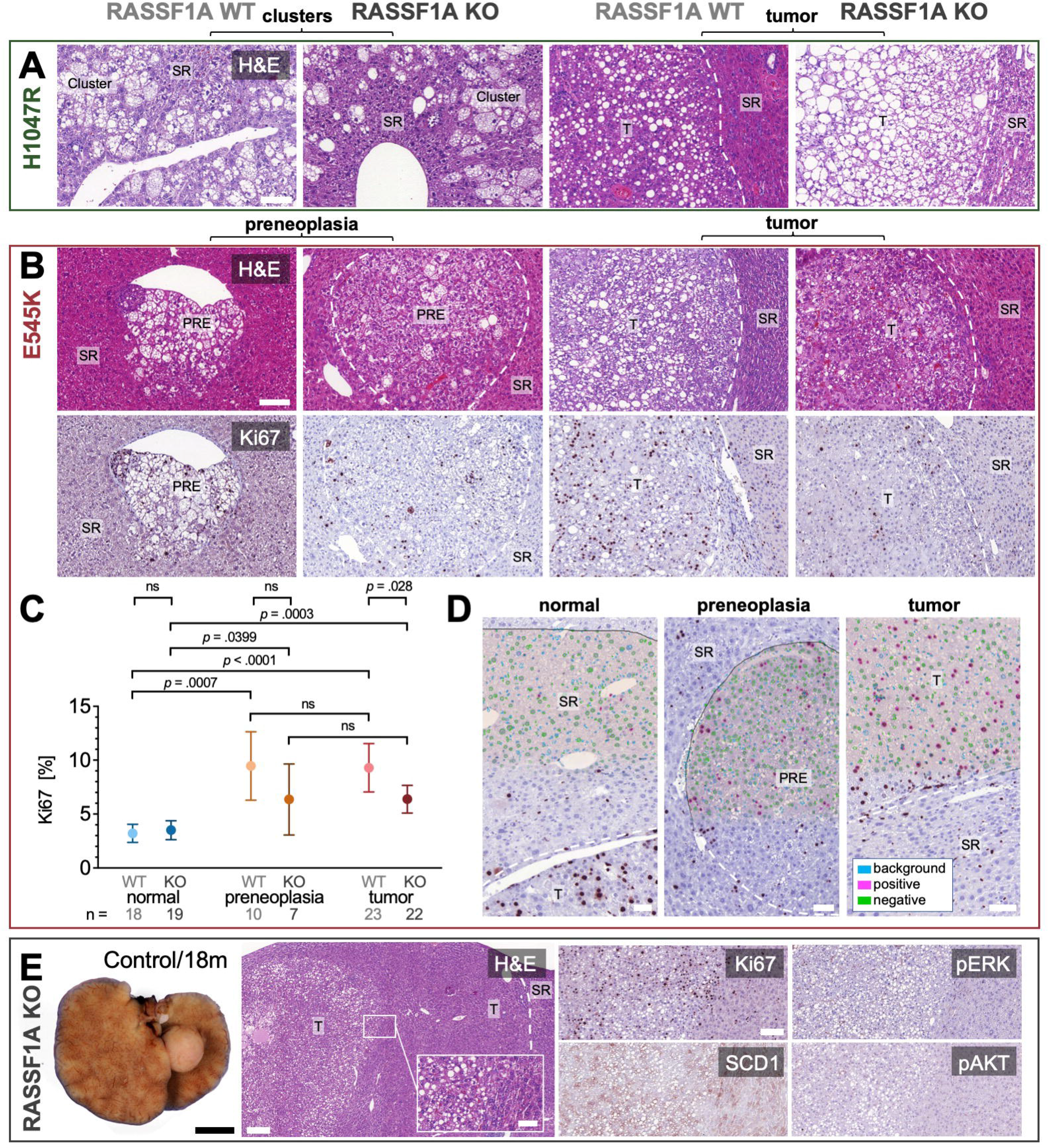
Histology and proliferation in PIK3CA dependent tumors induced in RASSF1A WT and KO mice. *(A)* Representative histological images of pericentral lipid-rich clusters and induced lipid-rich tumors resulting from PIK3CA H1047R injections in RASSF1A WT and KO mice. *Scale bar:* 100 µm. *(B)* Histological sections of PIK3CA E545K induced preneoplastic lesions and tumors in RASSF1A WT and KO mice with corresponding immunohistochemical staining for Ki67. *Scale bar:* 100 µm. *(B)* Quantification of Ki67 proliferation index in RASSF1A WT and KO mice in normal-appearing liver tissue, preneoplastic lesions, and tumors in PIK3CA E545K injected mice; n(mice, KO) = 22; n(mice, WT) = 24. Visualization as estimated mean with 95% confidence interval. *(D)* Screenshots obtained from DeePathology™STUDIO software exemplifying the region of interest selection (orange shading) and the detected nuclei with attribution of the properties background, positive or negative. Transparency gradient to original images without overlay from top to bottom. *Scale bars:* 50 µm. *(E)* Gross photograph of liver tumor spontaneously originating in an untreated RASSF1A mouse at 18 months experimental time point (corresponding to mouse age of 20 months, *left panel*). *Scale bar:* 0.5 cm. Exemplary histological images and immunohistochemical characterization of well-differentiated hepatocellular carcinoma transitioning into a lipid-rich central tumor component in a RASSF1A knockout PBS-injected control mouse at the experimental time point of 12 months (*right panels). Scale bars:* 200 µm (large panel); 50 µm (inset) and 100 µm (smaller panels). T, tumor; PRE, preneoplastic lesion; SR, surrounding tissue; pERK, phosphorylated extracellular-signal-regulated kinase; SCD1, Stearoyl-CoA desaturase; pAKT, phosphorylated RAC-alpha serine/threonine-protein kinase.

To ascertain the preneoplastic nature of the described lesions, we conducted a deep learning-based analysis using the software DeePathology^TM^ STUDIO after training the recognition of Ki67 positive and negative hepatocyte nuclei and the omission of background cells (such as lymphocytes and Kupffer cells) on manually selected regions of interest. A mean amount of 1650 cells were analyzed per category.

Mixed linear models (Figure 2*C*) demonstrated that preneoplastic lesions and tumors showed a significant increase in proliferation compared to normal-appearing tissue in *RASSF1A* WT and KO mice in PIK3CA E545K injected mice. At the same time, there was no significant difference in proliferation between preneoplastic lesions and tumors (Figure 2*C* and *D*). The proliferation was not significantly different between *RASSF1A* WT and KO mice in normal-appearing liver tissue and preneoplastic tissue (*p values* > .05). However, when comparing the proliferation in tumors between *RASSF1A* WT and KO, a statistically significant lower proliferation rate could be detected in *RASSF1A* KO mice (*p* = .0280). In addition, *RASSF1A* KO mice (m = 5.2, 95% CI = 3.7 – 6.7) compared to WT mice (m = 7.5, 95% CI = 6.1 – 8.9) unexpectedly showed a of reduced proliferation in tumors (*p* = .0331).

Finally, the two tumors spontaneously arising in *RASSF1A* KO mice in the combined control group were histologically examined. These tumors were extremely well-differentiated, pure HCC with a low proliferation rate demonstrated by Ki67 immunohistochemistry. Interestingly, one of these tumors featured a lipid-rich component with focally increased proliferation. This component displayed a hinted upregulation of SCD1. In contrast, phosphorylated/activated pERK and pAKT did not show an increased immunoreactivity (Figure 2*E*).

Altogether, the low-grade histology following a long latency confirms the limited oncogenic potential of *RASSF1A* inactivation alone. Proliferation showed a statistically significant tendency to be lower in PIK3CA E545K mutant tumors in *RASSF1A* KO mice than *RASSF1A* WT mice. Irrespective of *RASSF1A*, we could define the multistep nature of PIK3CA mutant forms induced hepatocarcinogenesis.

### 3.2. PIK3CA tumors display a strong upregulation of canonical effectors

Due to the absence of overt differences in tumorigenesis between *RASSF1A* WT and KO mice, we decided to focus on *RASSF1A* WT mice. To confirm the hepatocellular nature and the activity of PIK3CA effectors in the preneoplastic lesions and tumors developed in PIK3CA mutant mice, we performed immunohistochemical analyses. Immunohistochemistry demonstrated that pAKT, the primary downstream target of PIK3CA, was strongly upregulated in preneoplastic clusters and tumors. Similarly, pERK was markedly induced, implying the activation of the ERK-MAPK signaling pathway. Moreover, the master regulators of lipogenesis, SCD1 and FASN, were strongly positive both in E545K- and H1047R-injected cohorts. Intense positivity for carbamoyl phosphate synthetase I (CPS1) protein, a highly-specific hepatocyte marker, confirmed the hepatocellular nature of the neoplastic lesions. Compellingly, foci with cholangiocellular differentiation developed in PIK3CA E545K and H1047R mouse livers, as indicated by cytokeratin 7 (CK7) immunoreactivity. An overlap of CK7 with CPS1 was detected, which agrees with the hypothesis that the cholangiocellular components originate from transdifferentiation rather than from separate cholangiocarcinogenesis (Figure 3*A* and *B*) [50]. At the protein level, we verified the gradual upregulation of PIK3CA, p-AKT, p-SGK3, p-ERK1/2, p-STAT3, FASN, and SCD1 in preneoplastic lesions and tumors in PIK3CA E545K injected mice. Notably, Prostaglandin-endoperoxide synthase 2 (COX2), known to induce tumor-promoting inflammation through an increase in prostaglandin E2 synthesis [51], was also upregulated. Accordingly, eicosanoid metabolism is increasingly recognized as an essential mechanism of PIK3CA-mediated oncogenicity (Figure 3*C*) [52].

**Figure 3.**
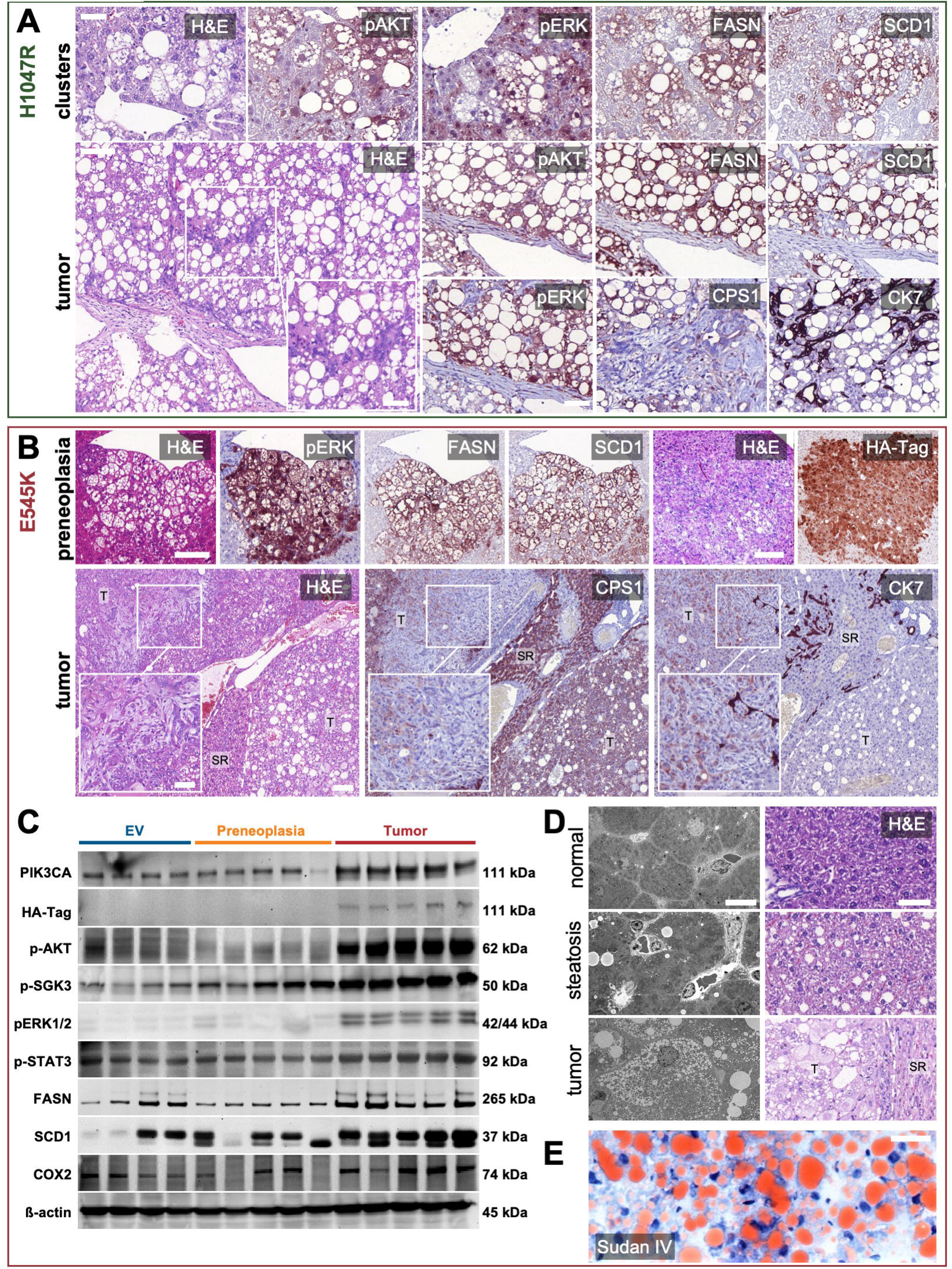
Activation of PIK3CA downstream effectors and the lipogenic phenotype in induced tumors. *(A)* Representative immunohistochemical analyses of preneoplastic clusters (*upper row*) and a tumor (*lower row*) generated by hydrodynamic tail vein injection of PIK3CA H1047R into wild-type mice. FASN and SCD1 represent master regulators of lipogenesis, while pAKT and pERK comprise downstream effectors of activated PIK3CA. Magnification inset highlighting focal cholangiocellular differentiation with the corresponding loss of CPS1 and positive staining for CK7. *Scale bars*: 50 µm (except for magnification inset: 100 µm). *(B)* Representative immunohistochemical analyses of a preneoplastic lesion (*upper row*) and a tumor (*lower row*) generated by hydrodynamic tail vein injection of PIK3CA E545K into wildtype mice. Tumor magnification inset illustrates spindle cell morphology with combined weak expression of CPS1 and CK7. *Scale bars*: first row and second row large panel 100 µm, magnification inset 50 µm. *(C)* Western blot analysis showing effective upregulation of PIK3CA and PIK3CA downstream effectors p-AKT, p-SGK3, p-ERK1/2, and p-STAT3, with increasing intensity in liver tissue containing preneoplastic lesions and tumors induced by PIK3CA E545K injections as compared to empty vehicle (EV) injections. Corresponding results for FASN, SCD1, and pro-inflammatory COX2. Loading control: β-actin. Molecular weights of observed bands are marked on the right. *(D)* Transmission electron micrographs of normal liver tissue of an untreated mouse at the experimental timepoint of 1 week (*upper panel*), an untreated mouse at the timepoint of 6 months with macrovesicular steatosis (*middle panel*), and a PIK3CA E545K induced tumor (*lower panel*) at a 9 months timepoint with diffuse microvesicular lipid inclusions. Adjacent matched H&E stained histological sections. *Scale bars*: 10 µm (electron micrographs), 50 µm (histological sections). *(E)* Histochemical Sudan IV staining of a PIK3CA E545K injected tumor highlighting cytoplasmic lipid droplets. *Scale bar:* 50 µm. T, tumor; SR, surrounding tissue; pERK1/2, phosphorylated extracellular-signal-regulated kinases 1/2; SCD1, Stearoyl-CoA desaturase 1; pAKT, phosphorylated RAC-alpha serine/threonine-protein kinase; CK7, cytokeratin 7; CPS1, Carbamoyl phosphate synthetase I; p-SGK3, phosphorylated serum/glucocorticoid regulated kinase family member 3; p-STAT3, phosphorylated Signal transducer and activator of transcription 3; COX2, Prostaglandin-endoperoxide synthase 2.

Next, we carried out transmission electron microscopy to evaluate the tumors on an ultrastructural level. In concordance with the histologically visible empty vacuoles, tumor cells showed abundant intracytoplasmic microvesicular lipid droplets. These differed markedly from an example of spontaneous liver steatosis in a control mouse, where only a few scattered, larger intracytoplasmic lipid droplets could be found. The extensive cytoplasmic accumulation of lipid vesicles underlines the profound reliance of these tumors on lipogenesis (Figure 3*D*). Subsequent Sudan IV histochemical staining on fresh frozen tissue confirmed the extensive presence of intracytoplasmic triglycerides in the tumors (Figure 3*E*).

Overall, we observed a marked upregulation of PIK3CA canonical downstream effectors. Massive lipogenesis, paralleled by robust upregulation of FASN and SCD1 lipogenic enzymes, was one of the earliest events in PIK3CA mediated hepatocarcinogenesis.

### 3.3. Identification of additional putative targets in PIK3CA driven hepatocarcinogenesis

Although lipogenesis is an established mediator of hepatocarcinogenesis, it remains challenging to target it selectively without significant systemic adverse effects due to its ubiquitous involvement in metabolism [53]. Therefore, the search needs to be widened to identify novel targets that can be pharmacologically manipulated. Thus, we performed gene expression microarray analyses of PIK3CA H1047R and E545K induced preneoplastic lesions and tumors (Figure 4*A*). *Lgals1*, which encodes Galectin-1, was among the few genes concomitantly upregulated in PIK3CA induced preneoplastic lesions and tumors compared with age-matched livers from empty vector-injected mice. We reached this conclusion based on the depicted Venn diagram [54] (Figure 4*B*). This finding was intriguing for several reasons. Galectin-1, being a 14 kDa beta-galactose specific binding protein [55], is known to be increased in numerous neoplasms, including primary hepatic tumors [56]. Moreover, Galectin-1 is a negative prognostic marker in HCC [57,58]. And finally, Galectin-1 has been linked to the PI3K-AKT-mTOR signaling, thereby enhancing epithelial-mesenchymal-transition (EMT) in HCC cell lines [59]. Given that Galectin-1 expression was already elevated in preneoplastic tissue and retained during tumor progression, it was selected for further investigation.

**Figure 4.**
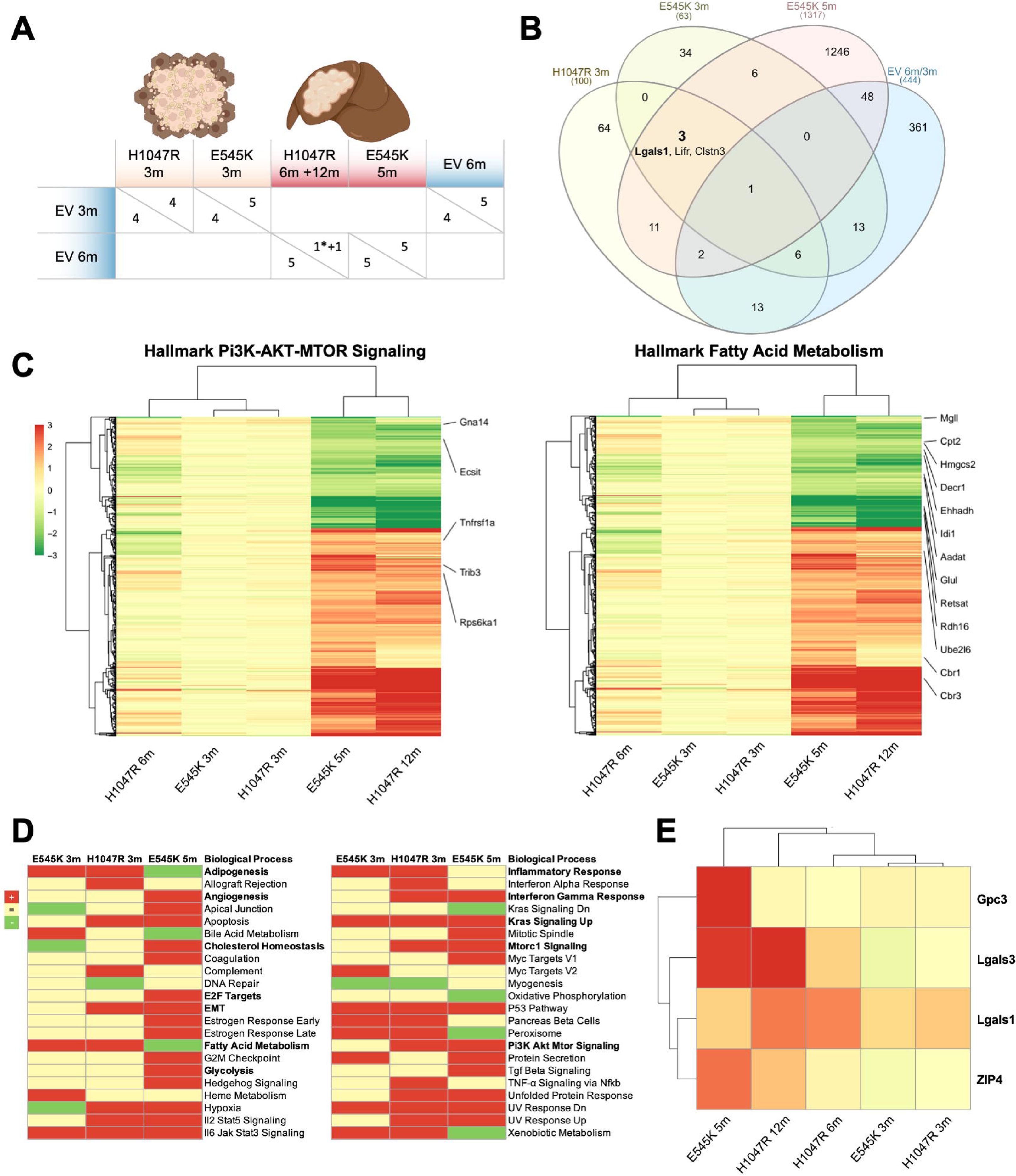
Metabolic signatures and identification of the novel effectors Galectin-1, Galectin-3, and ZIP4 in PIK3CA dependent hepatocarcinogenesis. *(A)* Overview of samples and comparison groups for cDNA microarray analysis. Preneoplastic lesions were compared to age-matched controls injected with the empty vector (EV). EV injected 3-month time points were contrasted to EV injected 6-months to rule out age-dependent effects. Only one tumor at 12 months was available from the H1047R injection arm. The asterisk indicates a PIK3CA H1047R induced confluent preneoplastic lesion analyzed at 6 months. *(B)* Venn Diagram of cDNA microarrays exhibiting overlap of regulated genes in the prespecified comparison groups. Selection criteria were Log1.5 change and *p* < .01 for H1047R 3 months, E545K 3 months and control 6 versus 3 months, Log2 fold-change and adj *p* < .05 for E545K 5 months. The numbers of overlapping and uniquely altered genes are given. The three genes conjointly called in the PIK3CA injection groups are mentioned, while Lgals1 encoding for Galectin-1 is written in bold. *(C)* Heatmap of differentially expressed genes with selection criteria |log2FC| > 1, adj. *p* < 0.05 and counts > 3 as determined in the E545K 5 months comparison group. Specified genes are comprised in the GSEA hallmark gene sets PI3K-AKT-MTOR signaling (*left panel*) and Fatty Acid Metabolism (*right panel*). Gradient scale color codes for Log2 fold-change. A cluster dendrogram is shown on the left and above. *(D)* GSEA was performed to identify significantly upregulated (red) or downregulated (green) biological processes in PIK3CA E545K 3 months, H1047R 3 months, and E545K 5 months injection groups. Particularly relevant processes implicated with hepatocarcinogenesis are reported in bold. *(E)* Target genes Gpc3 (Glypican-3), Lgals3 (Galectin-3), and ZIP4 fulfilled the selection criteria prementioned above in (*B)* and (*C)*. LGALS1 (Galectin-1) met the selection criteria described in (*B)*. Glypican-3 serves as an internal positive control as upregulation is expected in hepatocellular carcinoma. Target genes were manually chosen based on novelty, potential pharmacological targetability (Galectin-1 and Galectin-3), and early upregulation in preneoplastic lesions (Galectin-1). Presentation as a heatmap with identical color-coding as in Panel *B*.

In accordance with immunohistochemical results, induction of the PI3K-AKT-mTOR pathway was detected by the Hallmark Gene Set Enrichment Analyses (GSEA) [39]. The latter revealed an upregulation in preneoplastic lesions and/or tumors of pathways such as inflammatory response, interferon-gamma response, angiogenesis, cholesterol homeostasis, E2F targets, EMT, KRAS signaling, and Myc targets. Surprisingly, fatty acid metabolism was exclusively upregulated in preneoplastic lesions, suggesting a transition in the direction of increased independence from lipid metabolism in the later stages of carcinogenesis. Several genes in the Hallmark PI3K-AKT-mTOR gene set (upregulation of Tnfrsf1a, Trib3, and Rps6ka1; downregulation of Gna14, and Ecsit) and the Hallmark Fatty Acid Metabolism gene set (upregulation of Cbr3, Cbr1, Ube2l6; downregulation of Rdh16, Retsat, Glul, Aadat, Idi1, Ehhadh, Decr1, Hmgcs2, Cpt2, and Mgll) met the following criteria |log2FC| > 1, adj. *p* ≤ 0.05 and counts > 3 as determined in the E545K 5 months comparison group (Figure 4*C* and *D*).

Further analysis of the microarray data for targetable proteins with a cutoff of Log2 fold-change and adj *p* < .05 in the PIK3CA E545K injected group revealed *Lgals-3* and *ZIP4* as additional compelling targets (Figure 4*E*). Lgals-3 encodes Galectin-3, another member of the lectin family, implicated in the development of liver cirrhosis [60], immunosuppression in cancer [61], and with a negative prognostic value in HCC [62]. Zrt-Irt-like protein 4 (ZIP4) is a cellular zinc transporter [63,64], which is upregulated in HCC, where it represses apoptosis and increases migration [65]. Also, ZIP4 has been correlated with poor survival in HCC patients [66].

### 3.4. Galectin-1, ZIP4, and Galectin-3 are highly expressed in PIK3CA-induced lesions

To validate the gene expression microarray analysis results, the expression of Galectin-1, ZIP4, and Galectin-3 was investigated by Western blot analysis. All proteins were strongly upregulated in PIK3CA E545K induced tumors, and it was statistically significant for Galectin-1. Moreover, a tendency of upregulation occurred in preneoplastic lesions (Figure 5*A*). To corroborate these findings, we performed immunohistochemistry. Galectin-1 appeared robustly increased in tumors and preneoplastic lesions from PIK3CA mutant injected mice (Figure 5*B* and C).

**Figure 5.**
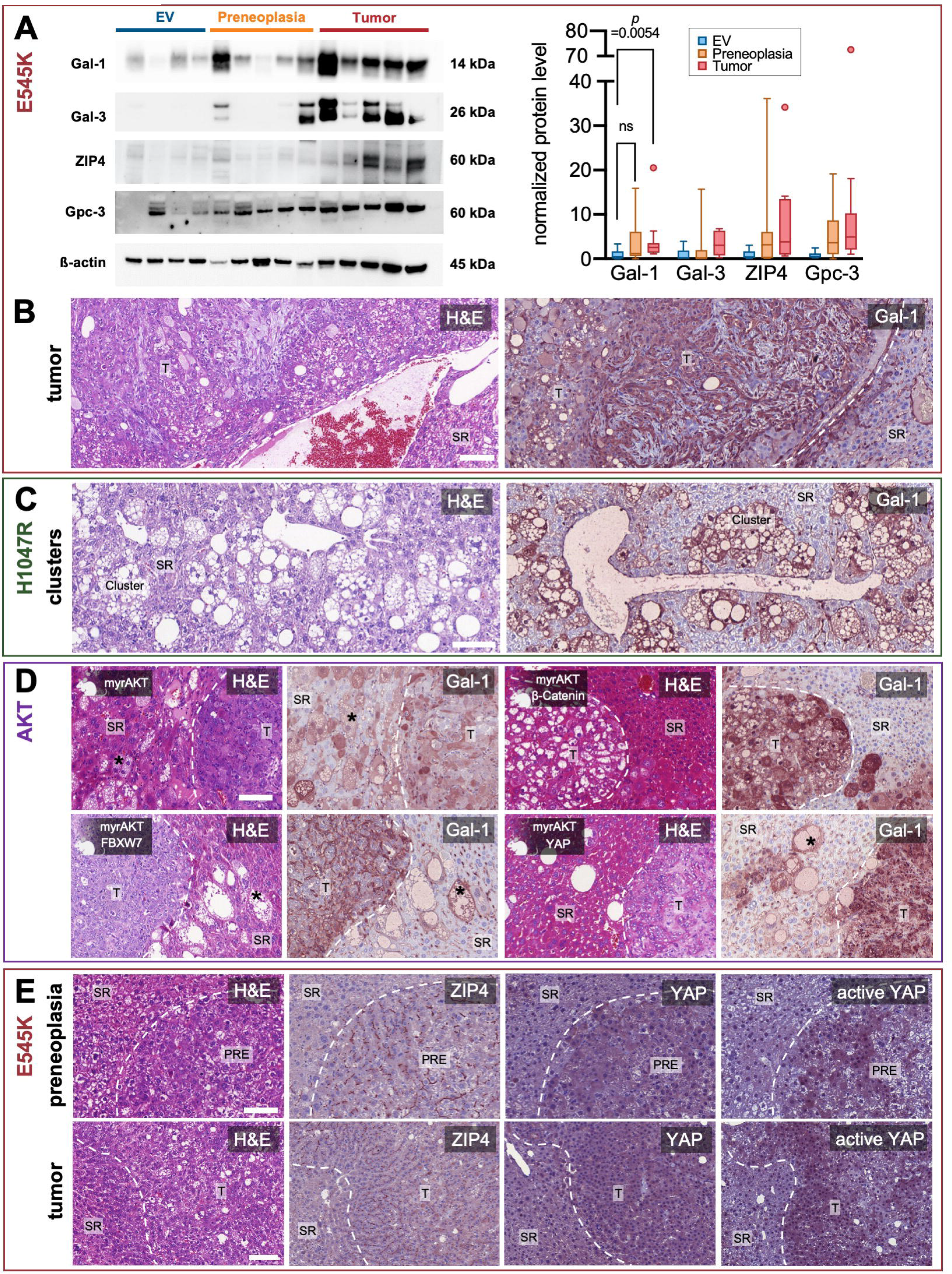
Validation of target proteins and early upregulation of Galectin-1 in precancerous lesions. *(A)* Western blot analysis demonstrating increased band intensities for Galectin-1 (Gal-1), Galectin-3 (Gal-3), ZIP4, as well as Glypican-3 (Gpc-3) in tumorous and in part also in precancerous liver tissue (*left panel*) in PIK3CA E545K injected mice, as compared to empty vehicle injections. β-actin was used as a loading control. Molecular weights of observed bands are marked on the right. Western blot quantification of band intensities normalized on corresponding β-actin signals (*right panel*) yielded a significant increase in Galectin-1 intensity of tumors versus preneoplastic lesions (*N* = 3 Western blot repeats). Kruskal-Wallis test was used. Tukey method box-and-whisker plots are displayed (ns = not significant; *p* > .05). Quantification of two Western blot repeats each is shown adjacently for the remaining target proteins. (*B*) Galectin-1 Immunohistochemistry of a PIK3CA E545K injection-induced tumor with spindle cell and steatotic components. *Scale bar*: 100 µm. (*C*) Galectin-1 Immunohistochemistry of preneoplastic pericentral clusters resulting from PIK3CA H1047R injection. *Scale bar*: 100 µm. (*D*) Galectin-1 overexpression in various mouse models generated by hydrodynamic tail vein injections of either activated AKT with myristoylation sequence alone (myrAKT) or in conjunction with β-Catenin, F-box/WD repeat-containing protein 7 (FBXW7), and Yes-associated protein 1 (YAP). Asterisks point out preneoplastic single cells. *Scale bar*: 100 µm. (*E*) ZIP4, YAP, and active YAP immunohistochemistry of a preneoplastic lesion (*upper row*) and a tumor (*lower row*) developing in a PIK3CA E545K injected mouse. Note the increased membranous expression of ZIP4 in contrast to surrounding liver tissue. *Scale bars*: 100 µm. T, tumor; PRE, preneoplasia; SR, surrounding tissue.

To investigate a potential link with the PIK3CA-AKT-mTOR pathway, we additionally examined different mouse models that were induced by the injection of myristoylated AKT either alone [67] or in combination with other genes, such as β-Catenin [68], F-box and WD repeat domain-containing 7 (FBXW7) [69], and YAP [70]. Notably, the latter two combinations induce cholangiocarcinogenesis [70]. In all mouse models, tumors exhibited strong immunoreactivity for Galectin-1 compared to the surrounding liver tissue. Remarkably, Galectin-1 positivity was also evident in isolated lipid-rich preneoplastic hepatocytes (Figure 5*D*).

Furthermore, we conducted immunohistochemical analyses of ZIP4 protein expression. Pronounced membranous immunolabeling for ZIP4 was detected in tumors and preneoplastic lesions of PIK3CA E545K injected mice. Interestingly, there was also a corresponding upregulation of the Hippo pathway effector YAP and active YAP (Figure 5*E*).

### 3.5. The Galectin inhibitor OTX008 synergizes with PI3K-inhibitors *in vitro*

In light of Galectin-1 overexpression in PIK3CA-AKT dependent tumors, we assessed the importance of Galectin-1 on HCC cell growth *in vitro*. Thus, we treated the PLC/PRF/5 and HLE HCC cell lines with the PI3K inhibitor Alpelisib, either alone or in association with the Galectin-1 inhibitor OTX008. Alpelisib has proven its clinical actionability in the large phase 3 trial SOLAR-1, which resulted in the drug’s approval by the Food and Drug Administration for hormone receptor-positive, human epidermal growth factor receptor 2–negative breast cancer [71]. In contrast, OTX008, a calixarene derivative designed to bind the Galectin-1 β-sheet conformation [72], has only undergone early clinical testing in the form of a dose-finding phase 1 study in patients with advanced solid tumors [73].

Combining the two drugs led to a significantly (by evaluation of 95% confidence intervals) [74] higher anti-growth properties than either of the two drugs alone. This effect was evident for combined concentrations of 20 and 30 µM in PLC/PRF/5 and 20 and 50 µM in HLE cell lines. Notably, OTX008 only displayed relevant anti-growth effects as a single drug in PLC/PRF/5 at the examined concentrations up to 50 µM (Figure 6*A*). To confirm the synergistic effect, the combination index was calculated based on the median-effect principle of the mass-action law [75]. A combination index <1 signifying synergism was attained for most combined concentrations in both PLC/PRF/5 and HLE cells (Figure 6*B*).

**Figure 6.**
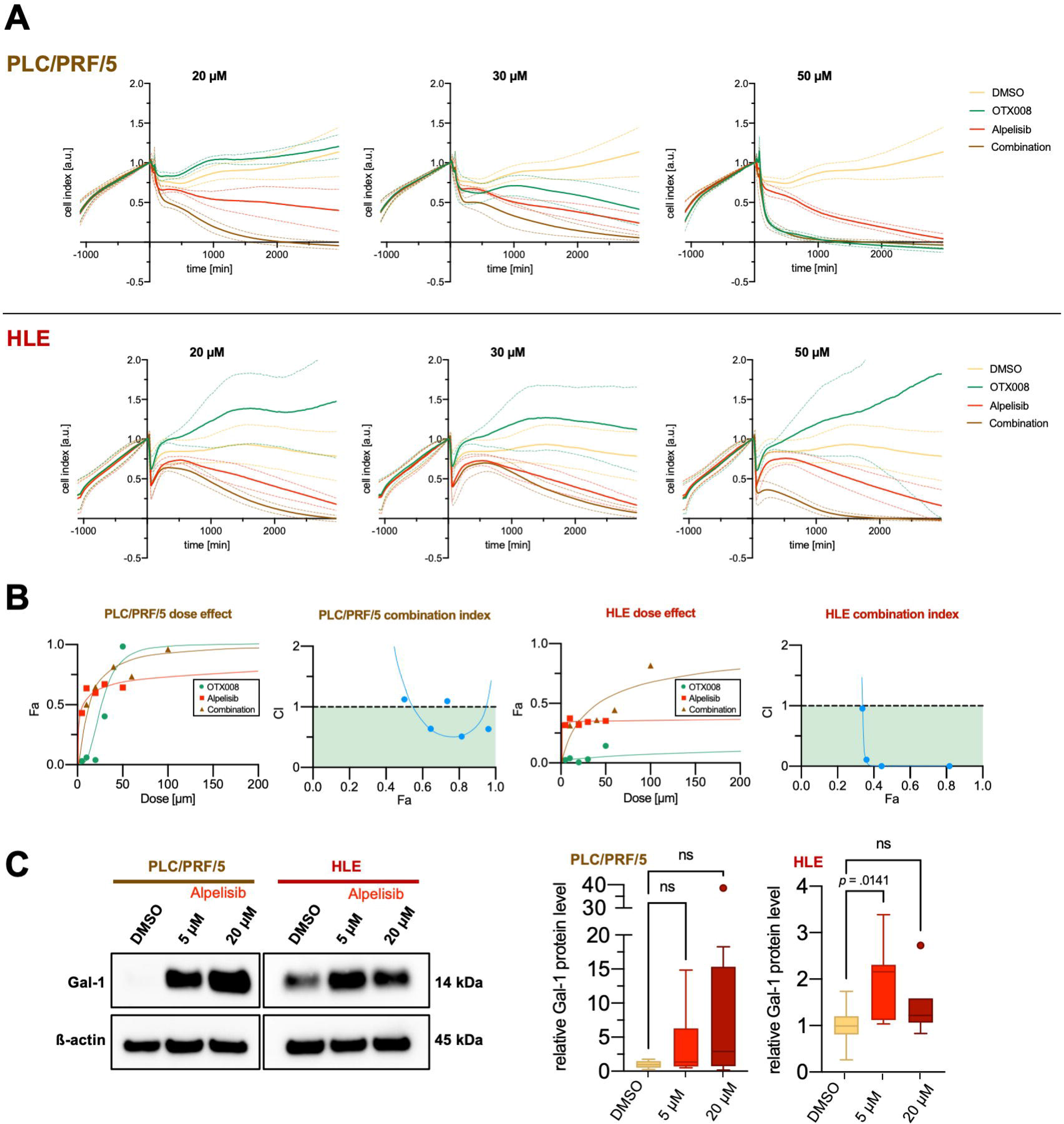
Strong synergism of Galectin-1 inhibitor OTX008 and PIK3CA-inhibitor Alpelisib in human hepatocellular carcinoma cell lines. *(A)* Time-dependent proliferation and cytotoxicity profiles generated by impedance measurements using an *xCELLigence®* device. PLC/PRF/5 cells are shown in the *upper row*, HLE in the *lower row*. Addition of matched DMSO, OTX008, Alpelisib at timepoint 0 in the outlined concentration either as a single or combined treatment after initial outgrowth for ∼24 hours. The cell index was normalized to the timepoint of drug addition. Solid lines signify mean and dashed lines 95% confidence interval. The illustrated data constitute 3 experimental repeats with 2-3 replicates acquired for each cell line and concentration. *(B)* CompuSyn®-software calculation results to determine drug synergism of OTX008 and Alpelisib in PLC/PRF/5 (*two left panels*) and HLE (*two right panels*). Values obtained by *xCELLigence®* cell viability analysis at 24 hours were employed, and the results of one representative experiment are shown. Dose-effect curves (Fa = fraction affected) and corresponding combination index plots are provided. A combination index (CI) < 1 is interpreted as synergism. Combination index (CI) values of < 1 (colored in green), 1, >1 can be interpreted as synergistic, additive, and antagonistic effects, respectively. (*C*) Exemplary Western blot illustrating a tendency of increased Galectin-1 levels in response to Alpelisib treatments in the HCC cell lines PLC/PRF/5 and HLE (*left panel*). A quantification of 3 treatment repeats with 2-3 replicates each was undertaken (*right panel*). Kruskal-Wallis test was used. ns, not significant.

To determine whether Alpelisib treatment influences Galectin-1 expression, we treated the two cell lines with 5 and 20 µM, respectively. Western blot analyses indicated an upregulation of Galectin-1 in response to Alpelisib treatment after 48 hours. However, this trend was only statistically significant (*p* = .0141) in the quantification at an Alpelisib concentration of 5 µM in the HLE cell line (Figure 6*C*). Although this finding contrasted our expectations, it might still explain the synergistic effects observed for combination treatments with OTX008, given that previous studies inferred a correlation between the antiproliferative effect of OTX008 and Galectin-1 expression [76].

To evaluate the generalizability of the observed synergy with OTX008 to other PI3K inhibitors, treatments of PLC/PRF/5 cells with 1 µM Taselisib, 1 µM Buparlisib, or 1 µM Alpelisib with or without the addition of 20 µM OTX008 were conducted for 72 hours. The mean excess over Bliss score was 0.1685 for Alpelisib, 0.1164 for Buparlisib, and 0.1548 for Taselisib in 3 internal replicates, indicating a similarly strong synergistic effect for all three of these compounds (Supplementary Figure 1).

Moreover, as a discovery approach for more potential interaction partners of OTX008 (at a concentration of 20 µM), we conducted a compound library screening including 315 approved drugs (at a concentration of 1 µM). All four PI3K inhibitors included in the screening demonstrated a synergistic effect in combination with OTX008. The excess over Bliss score was 0.0931 for Duvelisib, 0.0924 for Copanlisib, 0.0723 for Idelalisib, and 0.0189 for Buparlisib (Supplementary File 1). Using the threshold of viability from combination treatment < 0.6 and Z score of EOB > 1.5, several more putative synergistic interaction partners of OTX008 could be identified (Supplementary Figure 2 and Supplementary Table 1). Pacritinib, a Janus-Kinase 2 (JAK2)-inhibitor, emerged as the compound with the highest synergistic effect. On top of that, several other tyrosine kinase inhibitors, such as Axitinib, Apatinib, Crizotinib, Sorafenib, Crenolanib, and Nintedanib, and the two EGFR inhibitors Neratinib and Afatinib also reached the defined threshold. Thus, several approved drugs and drugs in advanced clinical trials could be identified as worthwhile candidates for combination treatments with OTX008, clearly warranting further investigations.

### 3.6. An interdependence with SCD1 links Galectin-1 to lipogenesis

In the search for a potential mechanism of cytotoxicity, we assessed whether Galectin-1 is associated with lipogenesis in HCC. A functional connection between Galectin-1 and the expression of lipogenic factors has recently been elucidated in livers of Galectin-1 knockout mice, where FASN, Acetyl-CoA carboxylase 1 (ACAC), and SCD1 were downregulated in animals fed a high-fat diet [77]. For this purpose, we performed Western blot analysis of SCD1 expression following Galectin-1 siRNA-mediated silencing (Figure 7*A*). As expected, Galectin-1 levels were downregulated in all tested HCC cell lines (*p* values < .05). Furthermore, decreased SCD1 levels in PLC/PRF/5, HLF, and Snu182 HCC cell lines subjected to Galectin-1 knockdown were observed. Quantitative analysis of band intensities supported this observation in PLC/PRF/5 and HLF and reached statistical significance (*p* values < .05), which was, however, not met in Snu182 (*p* = .34). In addition, we tested the effect of Galectin-1 downregulation on other lipogenic enzymes, including peroxisome proliferator-activated receptor gamma (PPARγ), ATP citrate synthase (ACLY), FASN, and ACAC. Among these, only PPARγ showed a statistically significant downregulation (*p* = .0173) upon Galectin-1 silencing in the PLC cell line (Figure 7*B*).

**Figure 7.**
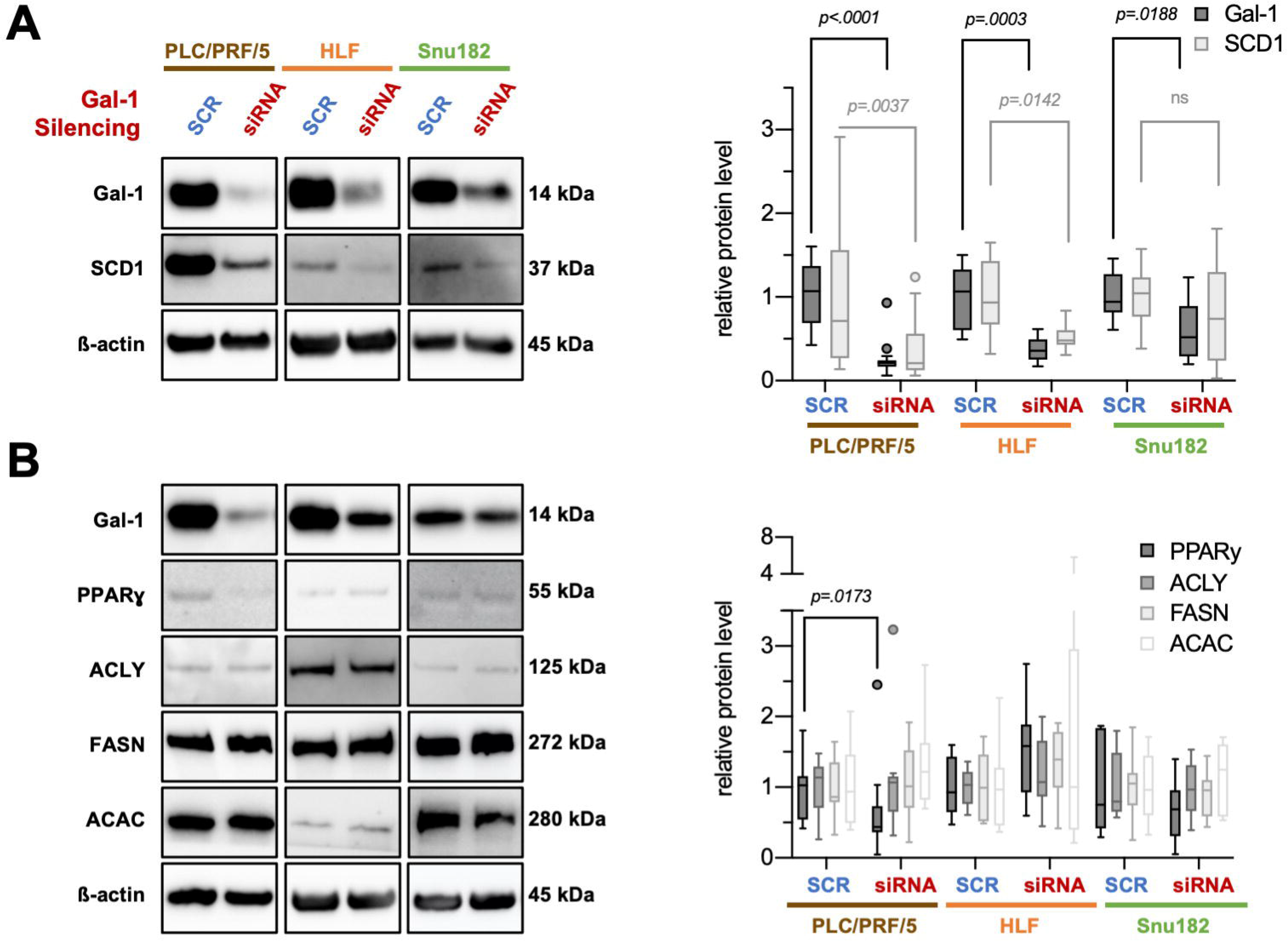
Interdependence of Galectin-1 and SCD-1 expression in human HCC cell lines suggests regulation of lipid synthesis. *(A)* siRNA-mediated Galectin-1 (Gal-1) silencing elicited a decrease in Stearoyl-CoA desaturase 1 (SCD1) protein levels in the human HCC cell lines PLC/PRF/5, HLF, and Snu182 as exemplified in Western blot analyses (*left panel*). Molecular weights of observed bands are noted. Quantification of yielded bands was performed (*right panel*). *(B)* In contrast, siRNA-mediated Gal-1 silencing did not yield a regulation of other important lipogenic enzymes, such as ATP citrate synthase (ACLY), Fatty acid synthase (FASN), and Acetyl-CoA carboxylase (ACAC). At the same time, a tendency for proliferator-activated receptor gamma (PPARɣ) reduction could be observed in PLC and Snu182 cell lines. n(PLC/PRF/5) = 3 cell culture experimental repeats with 3 cell culture replicates + 1 cell culture experimental repeat with 6 cell culture replicates. n(HLF, Snu182) = 3 cell culture experimental repeats with 3 cell culture replicates. Intensities were normalized to the mean of control for all tested proteins. Tukey method box-and-whisker plots are displayed *(right panels)*. Mann-Whitney tests were employed. ns, not significant. All pairwise comparisons in the *right panel* in *B* were not significant, except for the one displayed.

Taken together, we could deduce a dependence of SCD1, one of the master regulators of lipogenesis, on Galectin-1 expression in HCC.

## 4. Discussion

Both the PI3K-AKT-mTOR and the Ras-MAPK pathway have been commonly implicated in hepatocarcinogenesis [78,79]. This finding raised the question of possible interdependence and cooperativity between these signaling networks. RASSF1A as an inhibitor of Ras activity with a high methylation frequency in HCC [13] represented a promising candidate as an oncogenic signaling node, which could mediate the two pathways’ effects. We addressed the question of RASSF1A oncogenicity using a murine RASSF1A knockout model with hydrodynamic tail vein injections of the PIK3CA mutant forms H1047R and E545K. Surprisingly, we found that PIK3CA mutant forms elicited hepatocarcinogenesis irrespective of *RASSF1A* status, which precluded the hypothesized cooperativity.

Another unanticipated conclusion we could draw from these experiments was that PIK3CA mutant forms H1047R and E545K alone could induce hepatocarcinogenesis. Previous experiments suggested that combination partners such as activated NRAS or c-Met are necessary for PIK3CA to drive hepatocarcinogenesis [34]. Notably, PIK3CA E545K exhibited a much stronger oncogenic potential, giving rise to HCCs in most mice as early as three months after injection. PIK3CA H1047R, in contrast, only produced sporadic tumors after 12 months, which is opposite to findings in breast cancer [80]. The generation of this novel purely PIK3CA induced mouse model opens the door to further investigations into the PI3K-AKT-mTOR pathway in hepatocarcinogenesis. The availability of such a model is of high interest. First, because 4 to 6% of human HCCs harbor a PIK3CA mutation [7,32], which could be amenable to targeted therapies. Second, the PI3K-AKT-mTOR pathway activation often occurs in HCCs regardless of the PIK3CA mutational status [31]. Importantly, we could establish PIK3CA-mediated hepatocarcinogenesis as a multistep process including preneoplastic lesions, enabling the study of early predominant mechanisms. This finding paralleled observations that have been made in mice hydrodynamically injected with activated AKT [67]. When we examined the tumors induced in this model, we found an increase in the canonical effectors of PIK3CA, such as pAKT, pERK, p-SGK3, and p-STAT3. Moreover, COX2 was strongly induced, supporting a recent report implying the importance of eicosanoids in PIK3CA mutated tumors as they promote tumor-permissive inflammation [52].

In addition to these canonical targets, we revealed a pronounced increase of lipogenesis in tumors and even preneoplastic lesions, which was evident on several experimental layers: an increase in the level of the master regulators of lipogenesis SCD1 and FASN, an upregulation of lipid metabolism by gene expression analysis, and a lipid-rich phenotype to an extent, where tumor cells were packed with lipid droplets as seen by electron microscopy. *De novo* lipogenesis is a cancer hallmark. The synthesis of fatty acids provides tumor cells with metabolic intermediates essential for synthesizing the membrane lipid bilayer, energy storage in the form of triglycerides, and signaling molecules [53]. In a related murine HCC model, it has antecedently been demonstrated that the deletion of FASN abolishes AKT-induced carcinogenesis [81]. Therefore, inhibitors of FASN and other lipogenic enzymes can represent enticing therapeutic options, was it not for dose-limiting toxicity, which can arise in adipose and liver tissue [53]. Consequently, it is an important endeavor to widen the search for potential therapeutic targets that can influence lipogenesis.

Using gene expression analyses, we detected several candidates, which could serve as potential targets. Among them, we identified Galectin-1 as a protein already strongly upregulated in preneoplastic lesions, underlining its putative importance in hepatocarcinogenesis. A captivating finding was the interrelation of Galectin-1 expression and lipogenesis by regulating SCD1 protein levels. The incentive to analyze a potential link between Galectin-1 and lipogenesis came from a recent report showing the downregulation of FASN, ACC1, and SCD1 expression in adipose and hepatic tissues from Galectin-1 knockout mice [77]. This regulation could well play a critical role in hepatocarcinogenesis and warrants further experimental elucidation.

Intriguingly, Galectin-1 is accessible to pharmacological targeting, for instance, with the calixarene derivative OTX008, which has been tested in an early clinical trial [73]. In this study, the drug was administered to HCC cell lines. While there were limited effects when OTX008 was applied alone, a strong synergism could be detected in combination with the PIK3CA inhibitor Alpelisib. Indeed, previous reports directly put Galectin-1 in the context of PI3K-signaling [82]. In our hands, Alpelisib treatment resulted in an increase of Galectin-1 expression, which might sensitize cells to OTX008 treatment. In the phase 1 trial mentioned before, plasma concentrations “>1µM over 12h after administration” [73] were achieved. Given that the concentrations of OTX-008 were comparatively high in our experiments (∼ 20 µM), we must exert at least some caution in interpreting these findings. Of note, synergistic effects of OTX008 with either Sorafenib or Everolimus have been reported in HCC cell lines before [76], which further substantiates the interaction of Galectin-1 with the PI3K-AKT-mTOR pathway. In addition, multidrug drug screening revealed several more candidate drugs acting synergistically with OTX008, including JAK-, EGFR- and other tyrosine kinase inhibitors, highlighting its potential in combination therapies.

Finally, two additional upregulated targets identified in this study with potential therapeutic implications were Galectin-3 and ZIP4. Besides being repeatedly implicated in inflammation [83], Galectin-3 is also linked to lipid metabolism since FASN, proliferator-activator receptor gamma, and fatty acid-binding protein 4 were significantly reduced in Galectin-3 knockout mice [84]. ZIP4 emerged as a highly interesting target protein since we found a correlating upregulation of YAP and active YAP. ZIP4 and YAP have recently been demonstrated to be commonly responsible for EMT control in pancreatic cancer [85]. Further research is required to verify the context of these proteins in PIK3CA mediated carcinogenesis and determine the effect of ZIP4 on EMT in HCC.

Overall, the present study has determined the independence of *RASSF1A* knockout mediated and PIK3CA-driven hepatocarcinogenesis. We have established an HCC mouse model that is of high translational value concerning PI3K-targeted therapies. Furthermore, this model allowed us to confirm the paradigm of PIK3CA-driven carcinogenesis (in terms of canonical effectors of the pathway) and enhance our understanding of PIK3CA oncogenic properties. We ascertained the pivotal role of lipogenesis and discovered novel putative effectors, including Galectin-1, with therapeutic actionability (Figure 8). These findings pave the way for ensuing investigations, such as searching for further combination partners of Galectin-1 and the study of PIK3CA dependent oncogenicity in Galectin-1 knockout mice.

**Figure 8.**
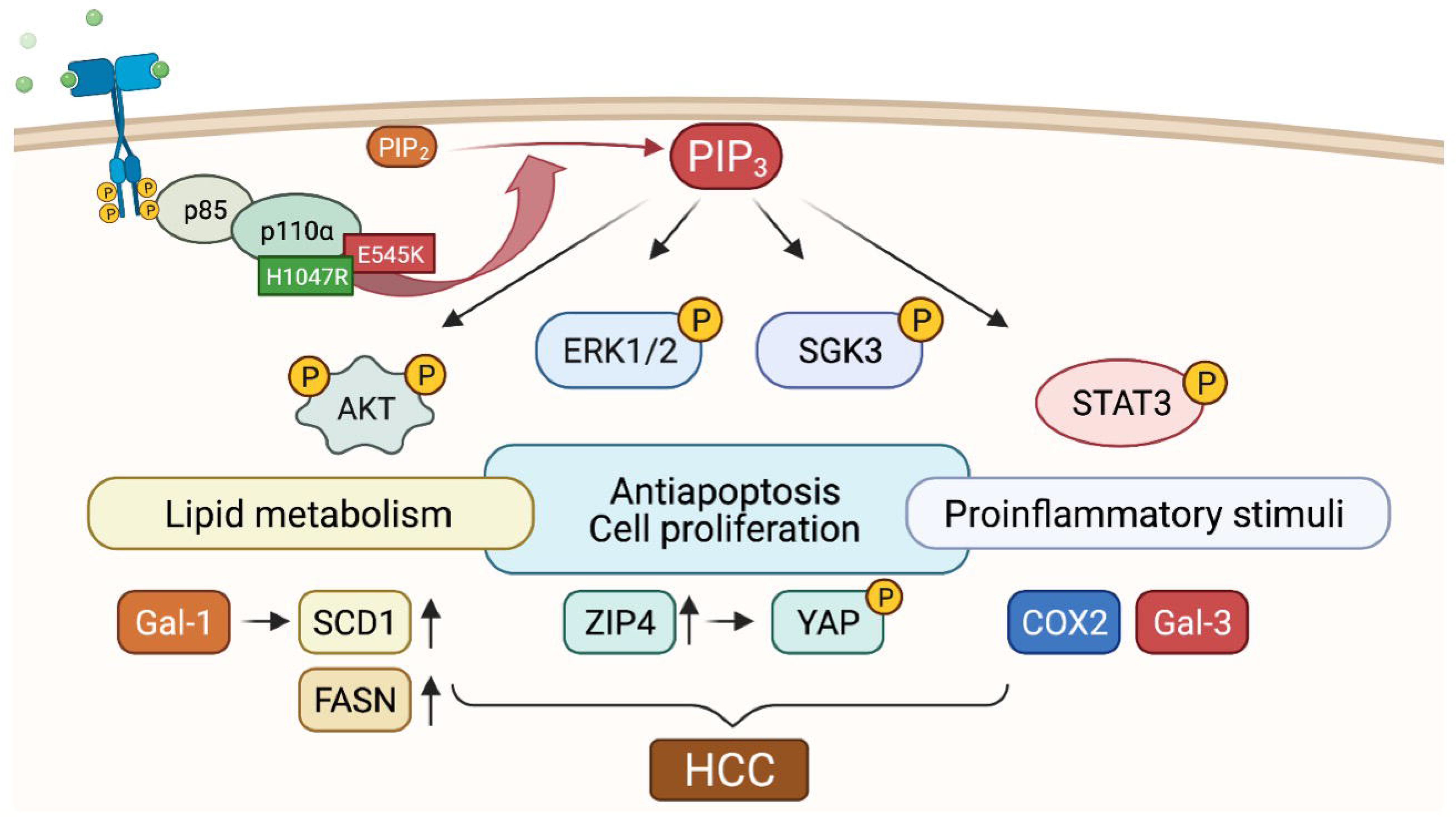
Graphical depiction of proposed PIK3CA induced hepatocarcinogenesis. H1047R and E545K mutations result in a receptor-independent constitutive activation of PIK3CA. In complex with the regulatory subunit p85, the conversion of phosphatidylinositol (4,5)-bisphosphate (PIP_2_) into phosphatidylinositol (3,4,5)-trisphosphate (PIP_3_) is catalyzed. PIP_3_ activates the downstream effectors RAC-alpha serine/threonine-protein kinase (AKT), Extracellular signal-regulated kinases 1/2 (ERK1/2), phosphorylated serum/glucocorticoid regulated kinase family member 3 (SGK3), and STAT3, either directly or via the mediation of phosphoinositide-dependent protein kinase 1. AKT has a pivotal role in the pathway by inducing antiapoptosis and cell proliferation. Cellular metabolism is veered toward lipid biosynthesis by increased expression of the master lipid regulators SCD1 and FASN. Galectin-1 upregulation acts in unison by eliciting the upregulation of SCD1 and could account for an accessible therapeutic target. Furthermore, the elevation of ZIP4 encompasses YAP activation, thereby potentially providing a bridge to the Hippo signaling pathway. Finally, in correlation with the macroscopic steatohepatitic phenotype, pro-inflammatory proteins such as COX2 and Galectin-3 (Gal-3) are stimulated. Cooperatively, these alterations induce the formation of hepatocellular carcinoma.

## 5. Conclusion

PIK3CA E545K and H1047R mutant forms confer oncogenic potential irrespective of RASSF1A *in vivo*. Induced HCCs showed a concomitant upregulation of lipogenesis and Galectin-1 expression. The regulation of SCD1 by Galectin-1 and the synergism of Alpelisib and OTX-008 suggest Galectin-1 as a potential therapeutic target in HCC.

## Abbreviations

ACAC: Acetyl-CoA carboxylase 1
ACLY: ATP citrate synthase
CK7: cytokeratin 7
COX2: Prostaglandin-endoperoxide synthase 2
CPS1: carbamoyl phosphate synthetase I
EOB: excess over the Bliss score
EMT: epithelial-mesenchymal-transition
EV: empty vector
FASN: Fatty acid synthase
FBXW7: F-box and WD repeat domain-containing 7
Gal-1: Galectin-1
Gal-3: Galectin-3
Gpc3: Glypican-3
GSEA: Gene Set Enrichment Analyses
HCC: hepatocellular carcinoma
JAK2: Janus-Kinase 2
KO: knockout
MAPK: Ras/Mitogen Activated Protein Kinase
myrAKT: AKT with myristoylation sequence
PI3K: phosphoinositide 3-kinase
PIK3CA: Phosphatidylinositol-4,5-bisphosphate 3-kinase, catalytic subunit alpha
PIP_2_: phosphatidylinositol (4,5)-bisphosphate
PIP_3_: phosphatidylinositol (3,4,5)-trisphosphate
PPARγ: peroxisome proliferator-activated receptor gamma
RASSF1A: Ras association domain-containing protein 1
SCD1: Stearoyl-CoA desaturase-1
SGK3: serum/glucocorticoid regulated kinase family member 3
STAT3: Signal transducer and activator of transcription 3
WT: wildtype
YAP: yes-associated protein 1
ZIP4: Zrt-Irt-like protein 4

---

**Supplementary Table 1:** Overview of drug targets with ≥ 1 hit

| Target | Hits | Non hits | All | Frequency |
| --- | --- | --- | --- | --- |
| COMT | 1.00 | 0.00 | 1.00 | 100.00 |
| CSF1 | 1.00 | 0.00 | 1.00 | 100.00 |
| CSK | 1.00 | 0.00 | 1.00 | 100.00 |
| FGFR4 | 1.00 | 0.00 | 1.00 | 100.00 |
| HIF1A | 1.00 | 0.00 | 1.00 | 100.00 |
| PLK4 | 1.00 | 0.00 | 1.00 | 100.00 |
| FGFR2 | 2.00 | 1.00 | 3.00 | 66.67 |
| CYP1A1 | 1.00 | 1.00 | 2.00 | 50.00 |
| CYP1B1 | 1.00 | 1.00 | 2.00 | 50.00 |
| FGFR3 | 2.00 | 2.00 | 4.00 | 50.00 |
| INSR | 1.00 | 1.00 | 2.00 | 50.00 |
| JAK3 | 1.00 | 1.00 | 2.00 | 50.00 |
| FLT3 | 4.00 | 6.00 | 10.00 | 40.00 |
| FGFR1 | 3.00 | 5.00 | 8.00 | 37.50 |
| KDR | 6.00 | 11.00 | 17.00 | 35.29 |
| DDR2 | 1.00 | 2.00 | 3.00 | 33.33 |
| ERBB2 | 2.00 | 4.00 | 6.00 | 33.33 |
| ERBB4 | 1.00 | 2.00 | 3.00 | 33.33 |
| FLT1 | 4.00 | 8.00 | 12.00 | 33.33 |
| MET | 1.00 | 2.00 | 3.00 | 33.33 |
| POLD1 | 1.00 | 2.00 | 3.00 | 33.33 |
| FLT4 | 4.00 | 9.00 | 13.00 | 30.77 |
| ALK | 1.00 | 3.00 | 4.00 | 25.00 |
| JAK1 | 1.00 | 3.00 | 4.00 | 25.00 |
| JAK2 | 1.00 | 3.00 | 4.00 | 25.00 |
| PDGFRA | 3.00 | 9.00 | 12.00 | 25.00 |
| PDGFRB | 4.00 | 12.00 | 16.00 | 25.00 |
| POLE | 1.00 | 3.00 | 4.00 | 25.00 |
| RRM2B | 1.00 | 3.00 | 4.00 | 25.00 |
| KIT | 4.00 | 13.00 | 17.00 | 23.53 |
| EGFR | 3.00 | 10.00 | 13.00 | 23.08 |
| CSF1R | 2.00 | 7.00 | 9.00 | 22.22 |
| MAP2K1 | 1.00 | 4.00 | 5.00 | 20.00 |
| POLA1 | 1.00 | 4.00 | 5.00 | 20.00 |
| RAF1 | 1.00 | 4.00 | 5.00 | 20.00 |
| RET | 2.00 | 8.00 | 10.00 | 20.00 |
| BRAF | 1.00 | 5.00 | 6.00 | 16.67 |
| RRM1 | 1.00 | 5.00 | 6.00 | 16.67 |
| RRM2 | 1.00 | 5.00 | 6.00 | 16.67 |
| CYP19A1 | 1.00 | 6.00 | 7.00 | 14.29 |
| TUBB | 1.00 | 9.00 | 10.00 | 10.00 |

**Supplementary Figure 1.**
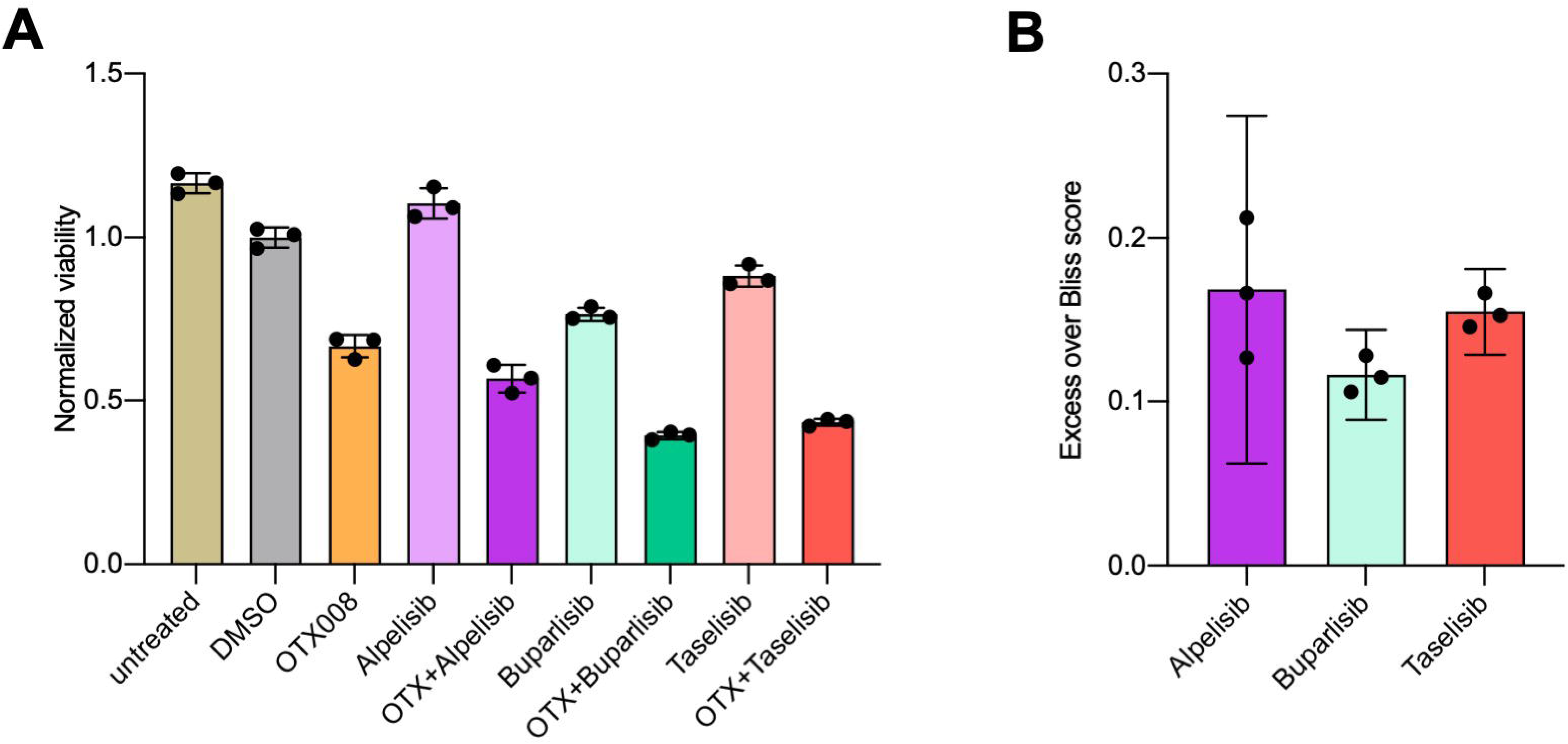
PLC/PRF/5 cells treated with indicated compounds in mono- or combination therapy for 72 hours. *(A)* Bar graph showing viability measured by ATP content normalized to mean of DMSO treated cells. B) Bar graph showing the excess over the Bliss independence model for each of the PI3K inhibitors combined with OTX008.

**Supplementary Figure 2.**
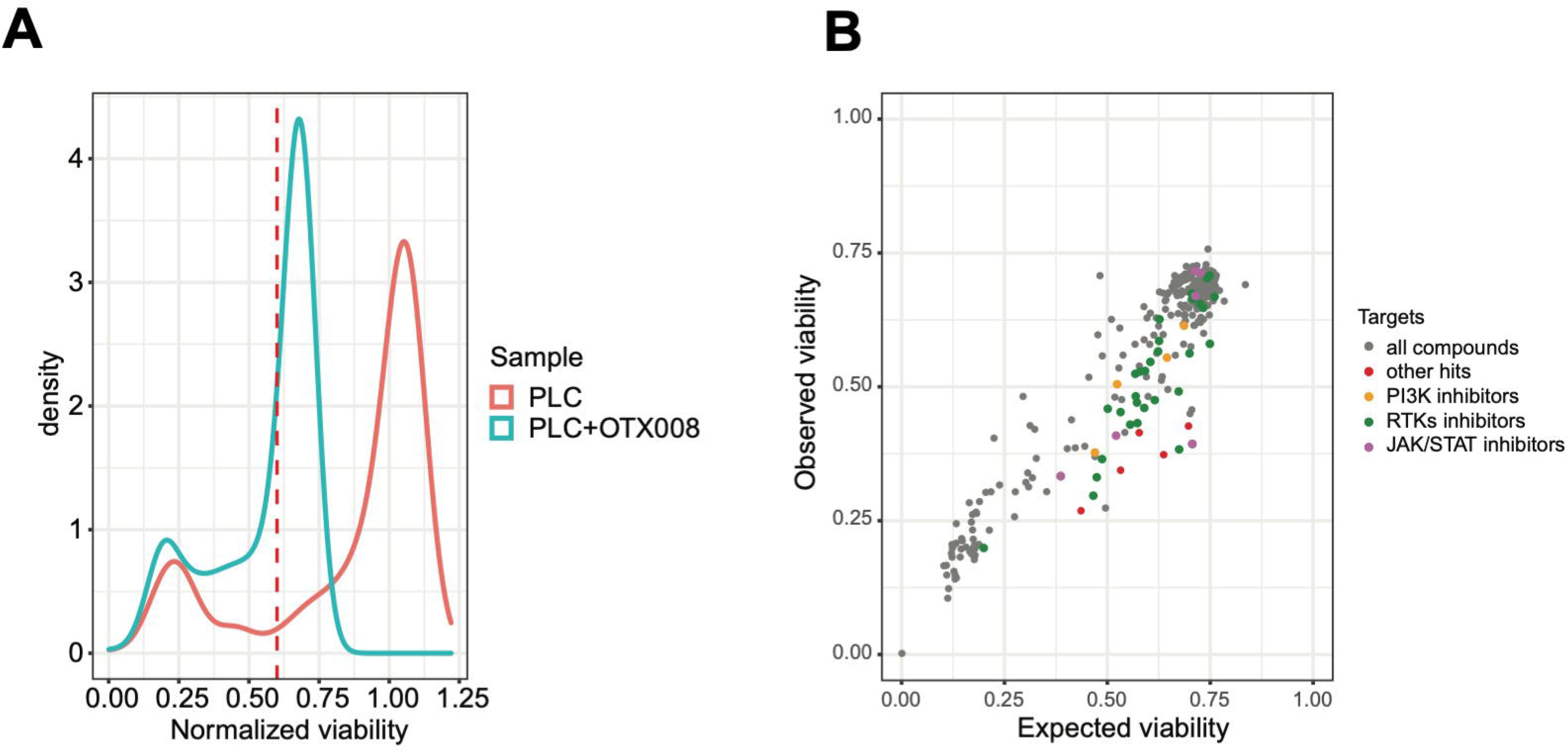
Drug screening of 315 approved anti-cancer compounds with and without OTX008. *(A)* Density plot of normalized viability data obtained from the screening of PLC/PRF/5 cells against a library of 315 approved anti-cancer compounds. Viability, measured as the number of Hoechst 33342 stained nuclei, is normalized to cells treated with DMSO only. The two conditions, with and without OTX008 1 µM are shown separately. The threshold of normalized viability = 0.6 is shown as a dashed red line (one of the parameters for a compound to be considered as a hit). *(B)* Scatter plot of observed normalized viability from the combinatory screening against the expected viability calculated from the measured normalized viability of single treatment according to the Bliss independence model. Compounds showing a normalized viability in the combination therapy < 0.6 and Z score of Excess over Bliss score of > 1.5 are defined as hits. Compounds targeting specific pathways are highlighted in respective colors: PI3K inhibitors in orange, tyrosine kinase inhibitors (RTKs inhibitors) in green and JAK/STAT inhibitors in violet.

## Data accessibility

The data that support the findings of this study are openly available in NCBI’s Gene Expression Omnibus and are accessible through https://www.ncbi.nlm.nih.gov/geo/query/acc.cgi?acc=GSE173963, GEO Series accession number GSE173963.

## Author Contributions

Conceptualization, K.E.., K.U., M.E., A.S., and D.C.; Methodology, K.U., K.E., M.E., A.S., and D.C..; Software, A.S., K.M., T.I., K.H., and A.C.; Validation, K.U., A.S., and D.C.; Formal Analysis, T.I., A.S., K.U., K.E., A.C.; Investigation, A.S., K.A., L.R., K.A., T.I., A.C., L.P; S.M.; Resources, M.E., X.C., F.D., A.T., C.B.; Data Curation, A.S., A.C., K.M.; Writing – Original Draft Preparation, A.S. and K.U.; Writing – Review & Editing, A.S. and D.C.; Visualization, A.S., T.I., and A.C.; Supervision, K.U., D.C., and M.E.; Project Administration, D.C.; Funding Acquisition, M.E.

## Acknowledgments

We thank our laboratory assistants, Ingrid Winkel and Manfred Meyer, for their invaluable support. Moreover, we thank Heiko Siegmund for his support with the acquisition of electron microscopic images and Ines Ziemack for mouse breeding. We acknowledge our use of the gene set enrichment analysis, GSEA software, and Molecular Signature Database (MSigDB) (Subramanian, Tamayo, et al. (2005), PNAS 102, 15545-15550, http://www.broad.mit.edu/gsea/). The graphical abstract, Figure 1*A*, Figure 4*A*, and Figure 8 were created with BioRender.com software. We acknowledge the use of DeePathology™ STUDIO for computer-aided Ki67 analysis. We thank Dr. Louise van der Weyden (Wellcome Trust Sanger Research Institute, Research Support Facility, Hinxton, Cambridge, CB10 1SA, UK) for providing a *RASSF1A* knockout mouse founder breeding pair.

## Supporting information

Additional supporting information may be found online in the Supporting Information section at the end of the article.

File S1. Combinatorial drug screening results for PLC/PRF/5 cells

